# Adverse structural and mechanical remodelling of main pulmonary artery in experimental pulmonary arterial hypertension is associated with impaired right ventricle-pulmonary artery coupling and function

**DOI:** 10.1101/2024.02.20.581269

**Authors:** Bahram Mirani, John D. Dauz, Kana Yazaki, Neda Latifi, J Paul Santerre, Michelle P. Bendeck, Craig A. Simmons, Mark K. Friedberg

**Affiliations:** Department of Mechanical & Industrial Engineering, University of Toronto, Toronto, Ontario, Canada, M5S 3G8; Institute of Biomedical Engineering, University of Toronto, Toronto, Ontario, Canada, M5S 3G9; Translational Biology and Engineering Program, Ted Rogers Centre for Heart Research, Toronto, Ontario, Canada, M5G 1M1; Labatt Family Heart Center, The Hospital for Sick Children, and Department of Paediatrics, University of Toronto, Toronto, Ontario, Canada; Department of Medical Engineering, University of South Florida, Tampa, Florida, USA, 33620; Faculty of Dentistry, University of Toronto, Toronto, Ontario M5G 1G6, Canada; Department of Laboratory Medicine and Pathobiology, University of Toronto, Toronto, Ontario, Canada

**Author notes:** Equal contribution (Bahram Mirani, John D. Dauz). Corresponding author: Mark K. Friedberg, MD, PhD, Labatt Family Heart Center, The Hospital for Sick Children, and Department of Paediatrics University of Toronto, 551 University Ave, Toronto, Ontario, Canada M5G1X8.

## Abstract

**Rationale:** Coupling between right ventricular function and the pulmonary vasculature determines outcomes in pulmonary arterial hypertension. The mechanics of the main pulmonary artery is an important but understudied determinant of right ventricular-pulmonary artery coupling.

**Objectives:** To investigate the histology and mechanics of the pulmonary artery in relationship to right ventricular remodeling, mechanics, hemodynamics and coupling in experimental pulmonary arterial hypertension.

**Methods:** In a sugen+hypoxia rat model of pulmonary arterial hypertension, right ventricular hemodynamics were assessed by conductance catheters. Active tension-strain curves were generated using echocardiography. Main pulmonary artery and right ventricle free-wall were harvested to determine their macro- and micro-structure, composition, and mechanical properties. Comprehensive multivariate analyses elucidated relationships between pulmonary artery and right ventricle mechanics, structure and coupling.

**Measurements and Main Results:** Pulmonary hypertensive main pulmonary arteries developed fibrosis relative to healthy controls, as did right ventricles, which also hypertrophied, with re-orientation of muscle fibres toward a tri-layer architecture reminiscent of normal left ventricular architecture. Increased glycosaminoglycan deposition and increased collagen-to-elastin ratio in the pulmonary artery; and increased collagen, as well as hypertrophy and reorganization of myofibers in the right ventricle, led to increased stiffness. This increase in stiffness was more pronounced in the longitudinal direction in the high- and low-strain regime for the pulmonary artery and right ventricle, respectively, causing increased mechanical anisotropy. Main pulmonary artery stiffening correlated significantly with right ventricular tissue mechanical remodelling and reduced systolic performance, cardiac output and right ventricle-pulmonary artery coupling.

**Conclusions:** Compositional, structural, and mechanical changes in the main pulmonary artery correlate with adverse right ventricular remodeling, mechanics, function and coupling in pulmonary arterial hypertension. Therefore, increasing mechanical compliance of the large pulmonary arteries may be an important and novel therapeutic strategy for mitigating right ventricular failure.

## Introduction

Pulmonary arterial hypertension (PAH) is a severe and often fatal disease characterized by progressive narrowing and stiffening of the pulmonary arteries (PA) due to pulmonary vascular remodeling involving endothelial and vascular smooth muscle cell (vSMC) proliferation, hypertrophy and hyperplasia, and fibrosis^1^. The resultant vascular proliferative lesions and fibrosis increase pulmonary vascular resistance (PVR) and reduce PA compliance^2^. While PVR is largely governed by the luminal size of the distal arteries, increased collagen deposition and loss of elastic components in both the proximal and distal PAs lead to increased pulmonary vascular impedance in PAH patients^3^.

Loss of main PA (mPA) compliance occurs early in PAH and contributes to disease progression^4^. Patients with only mild pulmonary vascular disease already have reduced mPA compliance, despite normal PA pressures and PVR^5^. Nonetheless, PVR is often used clinically in PAH to evaluate the RV afterload^6^ despite being a sub-optimal predictor of mortality^7^. Moreover, decreased mPA compliance correlates with the severity of PAH^5,3,8^ and with the degree of RV dysfunction, dilatation, and hypertrophy^9^, and may be a better predictor of mortality than PVR^3,4,6,10,11^. In this regard, the mPA and large branch PAs play a critical role in the transmission of pulsatile blood flow from the right ventricle (RV) to a near steady-state flow in the distal pulmonary vasculature via the Windkessel effect^12^.

RV function and response to the increased load imposed by PAH are major determinants of clinical outcomes^13–15^. The ability of the RV to efficiently maintain adequate cardiac output (CO) in the face of increased impedance and workload, i.e., RV-PA coupling, reflects RV adaptation to the increased load and correlates with clinical outcomes^16,17^. However, most emphasis has been placed on the distal pulmonary vasculature, while the histology and mechanics of the mPA remain incompletely characterized. Consequently, the relationship between mPA stiffness and RV remodelling, dysfunction, and RV-PA uncoupling is incompletely elucidated in the literature.

We aimed to characterize the structural, compositional, and mechanical remodelling of the mPA and RV, and to determine their relationship with RV hemodynamics and RV-PA coupling in a rat model of sugen plus hypoxia (SuHx) induced PAH. Histological and biochemical analyses and biaxial mechanical testing were utilized to characterize PA and RV compositional, architectural, and mechanical remodelling. Polarized light microscopy was used to elucidate regional changes in RV myocardial fibre microarchitecture. Comprehensive multivariate analyses were performed to determine correlations between PA and RV mechanics and correlations between mechanical remodelling and RV hemodynamics in SuHx animals compared to healthy controls.

## Methods

### Pulmonary arterial hypertension model

The experimental protocol was approved by the Hospital for Sick Children animal care committee in accordance with the Canadian Council on Animal Care (Protocol #: 1000059576). Rats were housed in a temperature-controlled (22±1℃) environment with 12-hr light/dark cycle and provided with standard commercial chow and acidified water *ad libitum*. Six-to seven-week-old male Sprague-Dawley rats (n=18), weighing 262±12g, were subcutaneously injected with a single dose of SU5416 (20 mg/kg). Rats were exposed to 10% O_2_ hypoxia for 3-weeks, followed by normoxia for an additional 3-weeks, as previously described^18^. Healthy age-matched rats were used as controls (n=19). Of the total number of rats, prior to the terminal experiment, RV function was assessed by echocardiography and cardiac catheterization in 8 healthy control and 6 SuHx rats. Following these procedures, the heart and great vessels were extirpated and the mPA isolated and excised. The heart and mPA were stored in ice-cold lactate Ringer’s solution (JB2324, Baxter, Missisauga, ON) prior to tissue analysis. Tissues harvested from the 8 healthy and 6 SuHx rats assessed by echocardiography and cardiac catheterization, and were dedicated to mechanical testing (for n=6 for each group; 2 healthy tissues failed during stretch cycles). From the remaining 11 healthy animals, 6 and 5 were dedicated to biochemical and histological analyses, respectively. The remaining 12 SuHx animals were divided into groups of n=6 for biochemical and histological analyses.

### Biochemical analysis

Quantitative biochemical assays were performed to measure mass of DNA, hydroxyproline (OH-proline), sulphated glycosaminoglycans (sGAGs), and elastin in healthy and SuHx mPA and RV tissues, cut with an equal planar area. Detailed methodology for these assays is provided in the Supplemental Materials.

### PA and RV free-wall histology

Excised tissues were embedded in optimal cutting temperature compound (Sakura, USA, cat# 4583), flash-frozen, sectioned into 7 µm slices at -20°C, and then stained with Movat’s pentachrome (STTARR Innovation Centre, University Health Network, Toronto) or picrosirius red (Abcam, Canada, cat# ab245887) according to the manufacturer’s protocol.

### Polarized light microscopy and image analysis

Fibrillar collagen content and organization in PSR stained sections of mPA and RV were assessed by quantitative polarization microscopy using the Abrio Imaging System (Cambridge Research and Instrumentation, USA) as previously described^19^. Because collagen fibers are oriented in many directions within a tissue, it is not possible to image all fibers at once with a single orientation of the polarizing filter. The Abrio system uses a circular polarizer with liquid crystal filters and a polarization algorithm to obtain orientation independent birefringent measurements of collagen fibers within the tissue. In these images, the brightness of each pixel is proportional to the birefringence retardance of the specimen. Therefore, the pixel intensity of the images reflects the amount of collagen, and the direction of fibers. The Abrio Imaging system was also used to measure the orientation and alignment of muscle fibres in the myocardium, taking advantage of the robust cytoskeletal contractile filaments in these cells which exhibit pronounced natural birefringence. Movat’s-pentachrome stained RV tissue sections were tile-imaged using a 40X lens, covering the complete cross-section of RV tissues. The images were pseudo-coloured by the microscope software to show fibre orientation with a colour spectrum. A MATLAB code was developed to assemble images, construct the entire tissue cross-section, and quantify fibre orientation based on pixel colour in the HSV colour space.

### Characterization of tissue passive mechanical properties

The biaxial tensile mechanical properties of mPA and RV free-wall tissues, harvested from the 8 healthy control and 6 SuHx rats, whose RV function was assessed by cardiac catheterization, were measured. The tissues were subjected to a series of loading cycles with a range of defined longitudinal/circumferential stretch ratios using a biaxial mechanical tester (BioTester 5000, CellScale, Canada), according to a previously developed method (**Figure S1A**)^20^. The experimental data collected from biaxial tensile tests were used to calculate Green strain, membrane tension, and material constants for a seven-parameter Fung model^21^. The material constants were used to describe mPA and RV tissue stiffness in the low- and high-strain regions of tension-strain curves in the longitudinal and circumferential directions. These calculations and the specific material constants used to describe tissue stiffness are detailed in the Supplemental Materials.

### Echocardiography

Rats were sedated using 2% isoflurane and placed on a temperature-controlled heated stage (Visualsonics Imaging Station, Toronto, ON). Excess fur was shaven and depilated. Echocardiography was performed using a 12-MHz phased-array probe (Vivid E9, GE Healthcare, Wauwatosa, WI). Two-dimensional short-axis views at the left ventricular (LV) papillary muscle level and apical 4-chamber views were acquired. The RV endocardial border was traced, and the RV curvature measured for active mechanical characterization (see below). Two-dimensional speckle-tracking echocardiography longitudinal strain curves were derived using RV apical 4-chamber views (EchoPAC, GE Healthcare, Wauwatosa, WI). RV endocardial borders were manually traced, and the region of interest was adjusted to the myocardial width, excluding trabeculations. Adequate tracking was visually verified. RV longitudinal strains were obtained using the lateral-free-wall basal, mid, and apical segments. RV circumferential strain was not adequately reliable and was excluded from the analysis.

### Characterization of RV active mechanical properties

Active tension in RV tissue was calculated by the Laplace equation using conductance catheter-pressures and RV curvature determined by echocardiography at end-systole and end-diastole:

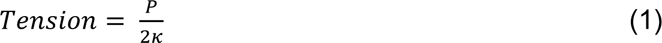

where P is pressure and *κ* is the mid-RV free-wall curvature. Active stiffness was calculated using:

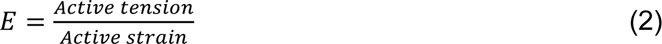

in which active strain was determined by speckle-tracking echocardiography.

### Hemodynamics

Cardiac catheterization was performed as previously described^22^. Rats were anesthetized with 2% isoflurane, intubated using a 14-gauge angiocatheter and ventilated with a volume-controlled ventilator (RoVent Jr., Kent Scientific Corp, Torrington, CT). Tidal volumes were adjusted for body weight. Thoracotomy was performed with rats in the supine position on a heating plate. While ensuring lungs were fully inflated, a 2-Fr conductance catheter (SPR-838, Millar Instruments, Inc., Houston, Texas) was advanced through the RV apex and pressure-volume (P-V) loops generated^23^. Data were acquired and analyzed using LabChart and MVPS Ultra (ADInstruments, Colorado Springs, CO). P-V loops used for analysis were obtained with a minimum of 5-10 beats with the ventilator off and the animal apneic. Measurements were obtained under steady-state conditions following inferior vena cava occlusion for preload reduction. Conductance signals were calibrated using fixed-volume cuvettes and stroke volume (SV) derived from echocardiography. RV parallel conductance was calculated by infusion of hypertonic saline (30% NaCl). RV functional parameters were calculated as previously reported^23^. P-V loop analysis yielded measures of RV systolic pressures and end-diastolic pressures (RVSP, EDP), stroke work (SW), time constant of monoexponential decay of RV pressure (tau, τ), and rate of RV pressure rise and decay (dP/dT_max_ and dP/dT_min_), arterial elastance (Ea), and SV. Analysis of P-V loops during preload reduction yielded end-systolic elastance (Ees) and preload recruitable stroke work (PRSW) as measures of RV contractility, and end-diastolic pressure-volume relations (EDPVR) as a measure of RV compliance. RV-PA coupling was calculated as a ratio of Ea and Ees (Ea/Ees).

### Statistics

Statistical analyses were performed using JMP Pro17. Data are presented as mean ± standard deviation (SD). Comparison of means between 2 groups with equal variance were analyzed by unpaired Student’s t-test. Comparisons of means with more than 2 groups with equal variance were analyzed using one-way analysis of variance (ANOVA) with Tukey’s post hoc test for pairwise comparisons. These comparisons were performed after confirming that the data were normally distributed. Wilcoxon/Kruskal-Wallis tests (rank sums) were used for comparisons of means for data with unequal variance. Pairwise correlation matrices were derived using multivariate analysis. Data were visualized using GraphPad Prism 9.

## Results

### Animal Characteristics

Body weight and surface areas (BSAs) of the 37 rats studied are presented in **Table 1**. The characteristics of rats used for RV function assessment and mechanical characterization are described in **Table 2**. SuHx animals had lower body weight values (p<0.0001), BSA (p≤0.0025), and heart-rate (p=0.048) when compared to controls.

**Table 1.**
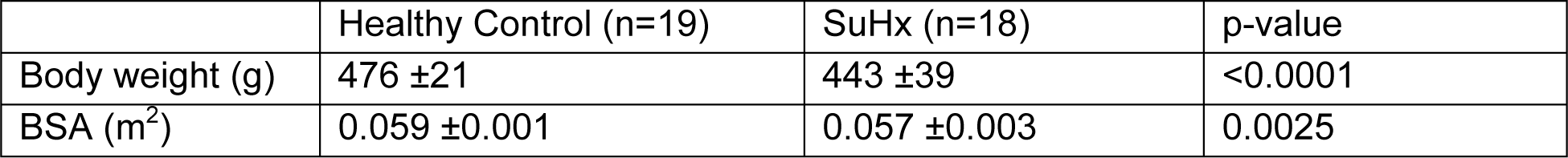
General characteristics of all rats used in the study. Data are presented as mean ± SD. SuHx: sugen hypoxia; BSA: body surface area.

|  | Healthy Control (n=19) | SuHx (n=18) | p-value |
| --- | --- | --- | --- |
| Body weight (g) | 476 ± 21 | 443 ± 39 | <0.0001 |
| BSA (m<sup>2</sup>) | 0.059 ± 0.001 | 0.057 ± 0.003 | 0.0025 |

**Table 2.** Characteristics of rats used for right ventricle function assessment and mechanical characterization. Data are presented as mean ±SD. SuHx: sugen hypoxia; HR: heart-rate; BSA: body surface area.

|  | Healthy Control (n=8) | SuHx (n=6) | p-value |
| --- | --- | --- | --- |
| HR (bpm) | 247 ± 25 | 219 ± 21 | 0.048 |
| Body weight (g) at cath | 465 ± 12 | 405 ± 34 | <0.0001 |
| BSA (m<sup>2</sup>) at cath | 0.0602 ± 0.001 | 0.054 ± 0.003 | 0.0001 |

### RV hemodynamics

RV hemodynamic parameters are described in **Table 3**. SuHx rats had significantly increased RV afterload as reflected by increased RV systolic pressures (RVSP), which were 87% of LV systolic pressures (LVSP), and increased Ea versus controls. SuHx rats had increased RV contractility as demonstrated by elevated Ees, PRSW, and maximum dP/dt. SuHx rats had reduced RV chamber compliance, as shown by the increase in the slope of the RV EDPVR and elevated RV EDP. SuHx rats also had impaired ventricular-arterial coupling ratio (Ea/Ees), reduced SV, and reduced CO versus controls.

**Table 3.** Right ventricle (RV) hemodynamics in healthy and sugen hypoxia (SuHx) rats. Data are presented as mean ± SD. RVSP: right ventricular systolic pressure; LVSP: left ventricular systolic pressure; EDP: end-diastolic pressure; SW: stroke work; CO: cardiac output; CI: cardiac index; SV: stroke volume; Ees: end-systolic elastance; EDPVR (end-diastolic pressure-volume relationship); PRSW: preload recruitable stroke work; Ea: arterial elastance.

|  | Healthy Control (n=8) | SuHx (n=6) | p-value |
| --- | --- | --- | --- |
| RVSP (mmHg) | 27 ± 4 | 69 ± 9.6 | <0.0001 |
| RVSP/LVSP (%) | 35 ± 3 | 87 ± 16 | <0.0001 |
| RV EDP (mmHg) | 5.9 ± 1.1 | 7.6 ± 1.8 | 0.0465 |
| SW (mmHg*μL) | 6586 ± 1412 | 12250 ± 2054 | <0.0001 |
| CO (mL/min) | 90.3 ± 14.4 | 59 ± 16.1 | 0.0024 |
| CI (mL/min/m<sup>2</sup>) | 1501 ± 253 | 1090 ± 269 | 0.0127 |
| SV (mL) | 0.36 ± 0.03 | 0.27 ± 0.05 | 0.0005 |
| dP/dT<sub>max</sub> | 961 ± 193 | 2275 ± 303 | <0.0001 |
| dP/dT<sub>min</sub> | -674 ± 175 | -1424 ± 221 | <0.0001 |
| Tau (s) | 0.035 | 0.028 | 0.37 |
| Ees (mmHg/μL) | 0.029 ± 0.009 | 0.495 ± 0.212 | <0.0001 |
| EDPVR (mmHg/μL) | 0.003181 ± 0.0016 | 0.02874 ± 0.025 | 0.0133 |
| PRSW | 12 ± 5 | 58 ± 12 | <0.0001 |
| Ea (mmHg/μL) | 0.073 ± 0.01 | 0.272 ± 0.087 | <0.0001 |
| Ea/Ees | 2.78 ± 1.25 | 0.60 ± 0.17 | <0.0001 |

### PA structural remodelling

In SuHx rats, the mPA was enlarged (**Figure 1A**) with increased diastolic diameter of 1.3-fold (p=0.0016), increased relaxed (passive) length of 1.4-fold (p<0.0001), and wall thickening of 3.2-fold (p=0.0040; **Figure 1B**). Histological analysis using Movat’s pentachrome (**Figure 1C**) and PSR (**Figure 1D**) staining on circumferential and longitudinal sections of mPA tissue showed increased PA adventitia collagen deposition (yellow in Movat’s and red in PSR) in SuHx rat group. Polarized light microscopy analysis of PSR-stained sections showed higher retardance levels in SuHx mPA tissues, reflecting their more organized, fibrillar collagen structure compared to healthy tissues (**Figure 1E**). Derived from these images, SuHx mPA showed increased mean pixel intensity (indicative of collagen fibrillar organization and content per tissue cross-sectional area) in the longitudinal direction and increased total intensity (indicative of collagen fibrillar organization and total content) in both longitudinal and circumferential directions (**Figure 1F**).

**Figure 1.**
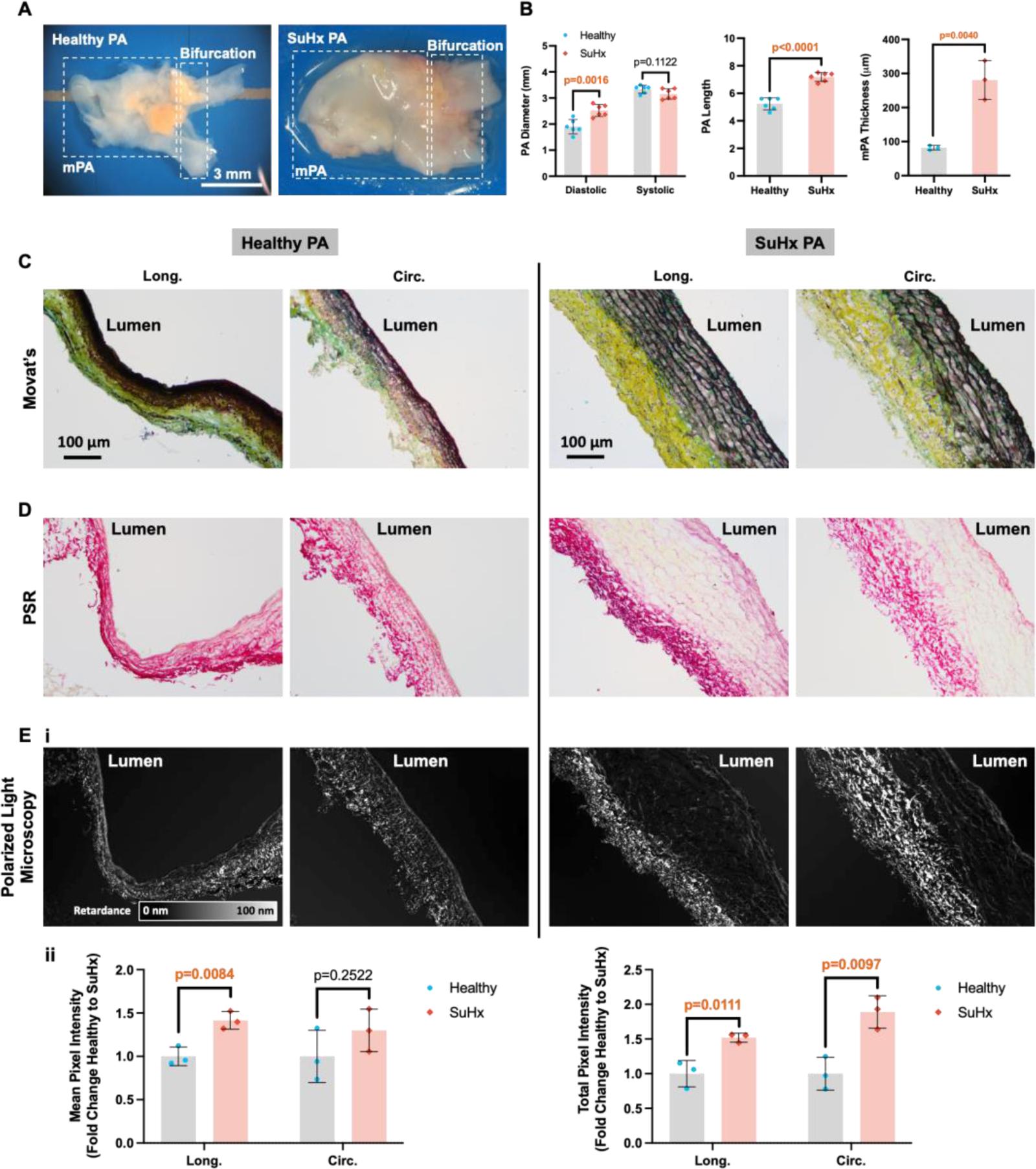
Pulmonary artery (PA) tissue structure. **A)** Images of PA tissues harvested from healthy and sugen hypoxia (SuHx) rats. **B)** Diameter, length, and thickness of SuHx and healthy PA tissues. Data are presented as mean ± SD with n=3 healthy and n=3 SuHx rats. **C)** Movats’s pentachrome staining images of healthy and SuHx PA tissues in the longitudinal (Long.) and circumferential (Circ.) directions. **D)** Picrosirius (PSR) staining images of healthy and SuHx PA tissues. **E)** Polarized light micrographs of healthy and SuHx PA tissues performed on PSR-stained sections (i). Quantification of pixel intensity based on polarized light micrographs for healthy and SuHx PA (ii). Total intensity represents the product of mean intensity and tissue thickness.

### RV structural remodelling

The RV myocardium was significantly thickened in SuHx rats by 2.3-fold (p<0.0001; **Figures 2A,B**). Movat’s pentachrome staining showed relatively uniform distribution of collagen in the healthy control RV tissues, in contrast to increased accumulation of interstitial collagen fibres in SuHx RV tissues (**Figure 2C**). This observation was confirmed by PSR staining and polarized light microscopy, where collagen appeared more abundant, organized, and fibrillar in the SuHx tissues with localized deposition (**Figure 2D**) and higher retardance (**Figure 2E**). These findings were more prominent in the longitudinal compared to the circumferential direction (**Figure 2E-ii**). Polarized microscopy on Movat’s pentachrome stained sections of RV tissue showed a transformation in muscle fibre alignment and organization from healthy control RV tissues to PAH (**Figure 3**). Fibres oriented longitudinally or circumferentially were found in all layers of healthy RV tissue from epicardium and endocardium (**Figure 3A**). In contrast, SuHx RV tissues acquired a distinct trilayer muscle fibre organization with fibres in the epicardium and endocardium predominantly oriented longitudinally (**Figure 3B-i**) and those in the mid-wall oriented circumferentially (**Figure 3B-ii**). This transformation in fibre organization was seen in 5 out of 6 SuHx rats used in this analysis (**Figure S2**). Overall, in SuHx RV tissues, muscle fibres were aligned predominantly in the longitudinal direction (**Figure 3C**).

**Figure 2.**
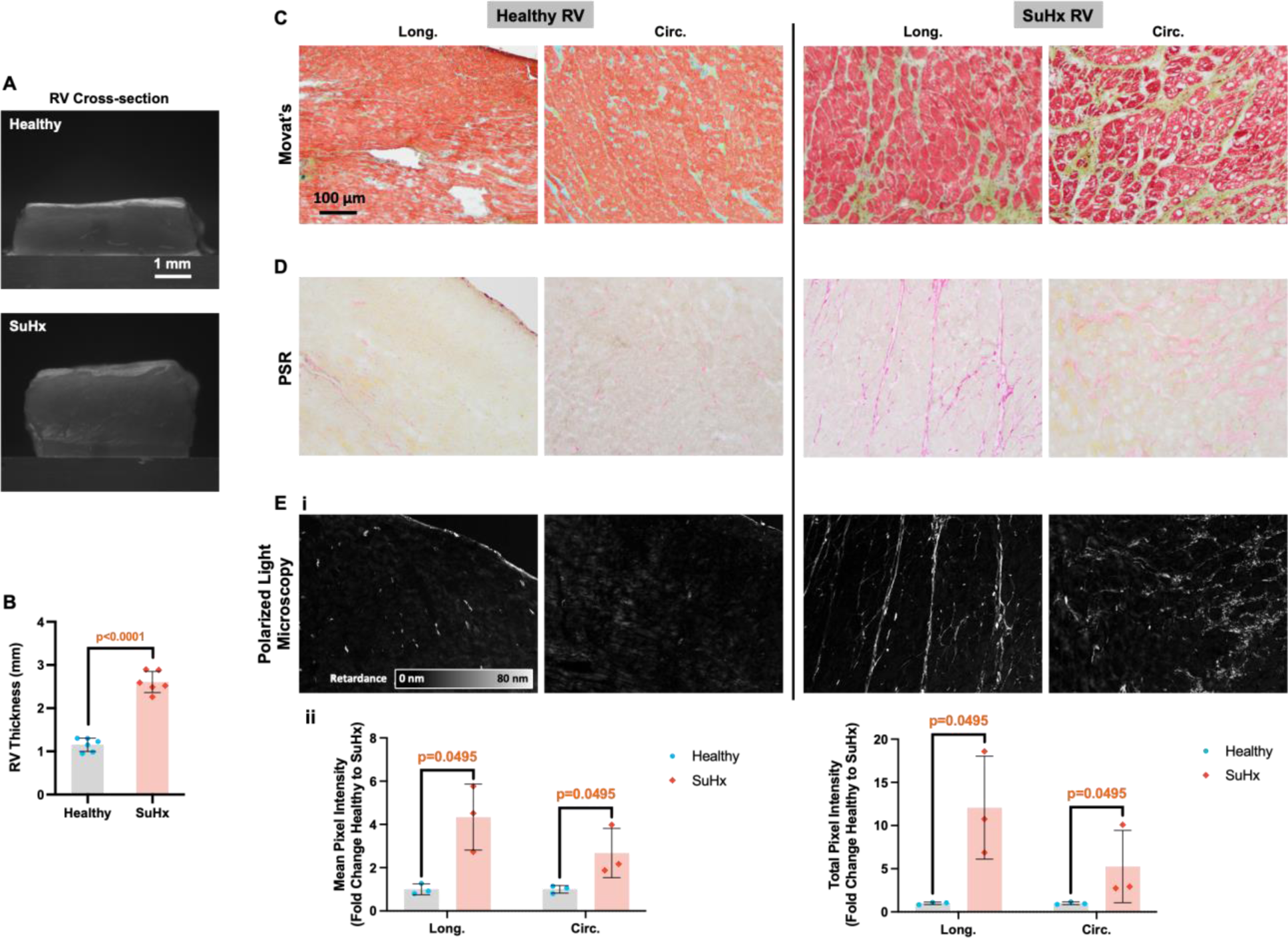
Right ventricle (RV) tissue structure. **A)** Microscopic images showing the cross-section of healthy and sugen hypoxia (SuHx) RV tissue. Images were taken using a 12X zoom Navitar® system. **B)** Thickness of healthy and SuHx RV tissues. Data are presented as mean ± SD with n=6 healthy and n=6 SuHx rats. **C)** Movat’s pentachrome staining images of healthy and SuHx RV tissues in the longitudinal (Long.) and circumferential (Circ.) directions. **D)** Picrosirius (PSR) staining images of healthy and SuHx RV tissues. **E)** Polarized light micrographs of healthy and SuHx RV tissues performed on PSR-stained sections (i). Quantification of pixel intensity based on polarized light micrographs for healthy and SuHx RV (ii). Total intensity represents the product of mean intensity and tissue tickness.

**Figure 3.**
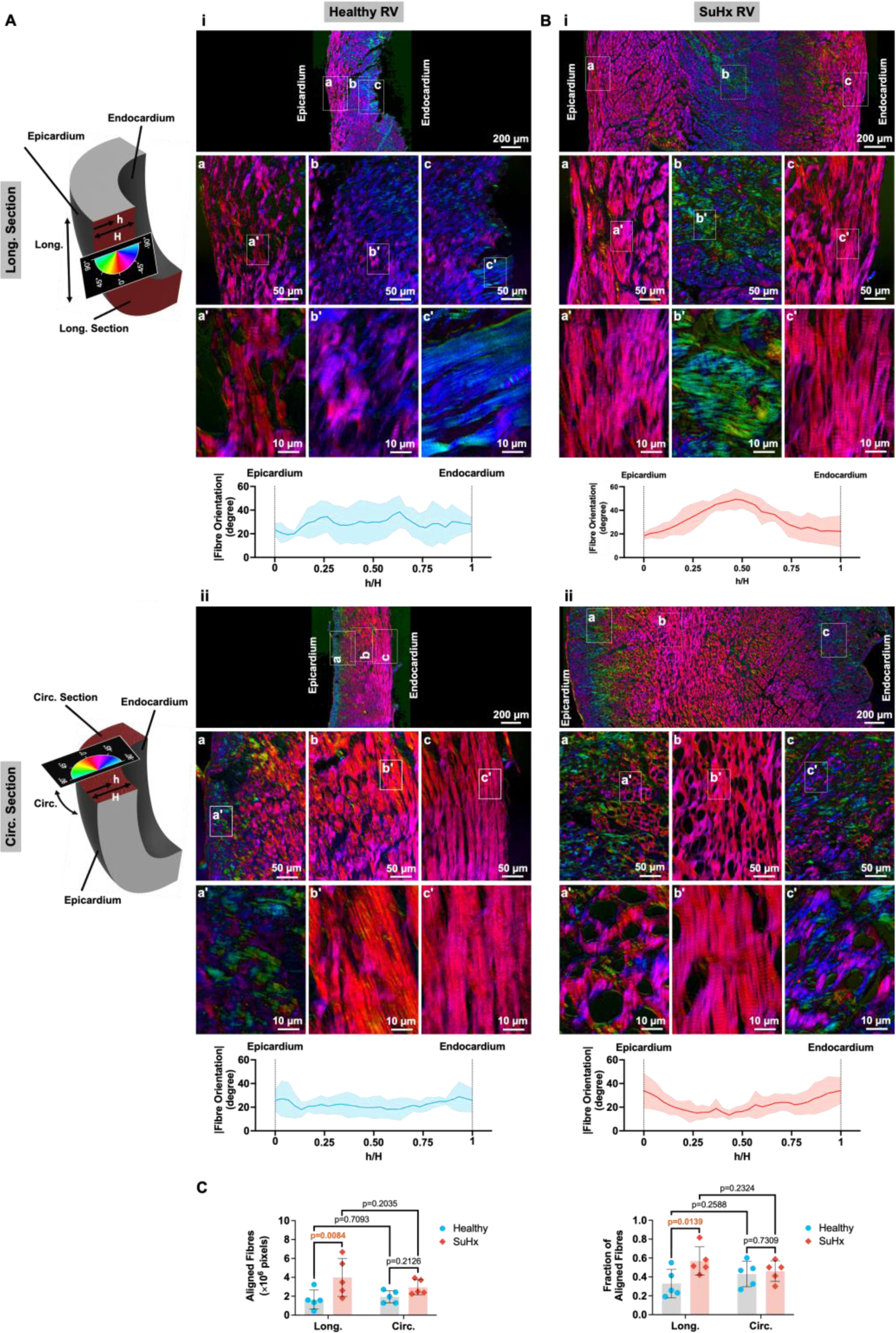
Muscle fibre organization in the right ventricle (RV) tissue. **A)** Polarized light microscopic imaging was performed on Movat’s pentachrome stained sections of healthy tissues in the longitudinal (Long.) (i) and circumferential (Circ.) (ii) direction. The images were pseudo-coloured to show and quantify fibre orientation. Shaded region of the graphs shows SD. H represents myocardial thickness and h represents a length variable starting from epicardium to endocardium (h/H is equal to 0 and 1 at epicardium and endocardium, respectively). **B)** Pseudo-coloured polarized light microscopic images showing the orientation of fibres in the longitudinal (i) and circumferential (ii) direction of sugen hypoxia (SuHX) RV tissue. Shaded region of the graphs shows SD. **C)** Fibre orientation quantified as total (based on pixel count where deviation from Long. or Circ. direction is within ±15°) and fraction in the healthy and SuHx RV tissues. Data are presented as mean ± SD with n=5 healthy and n=5 SuHx rats.

### mPA and RV compositional remodelling

mPA biochemical analysis showed increased absolute total mass of DNA, collagen (via quantification of OH-proline), sGAGs, and elastin in SuHx (**Figure 4A**). However, when normalized to tissue dry weight, masses were either similar between the two groups, or higher in healthy controls. The ratio of PA collagen/elastin content was considerably higher in SuHx (3.4±0.5) versus healthy control (1.6±0.7) rats (p=0.0007; **Figure 4B**). Similarly, RV tissues in the SuHx group showed increased total DNA, collagen, and sGAG mass relative to healthy controls (**Figure 4C**). However, when normalized to tissue dry weight, no significant differences were observed in DNA or collagen content, and sGAG content was reduced in SuHx RV tissue.

**Figure 4.**
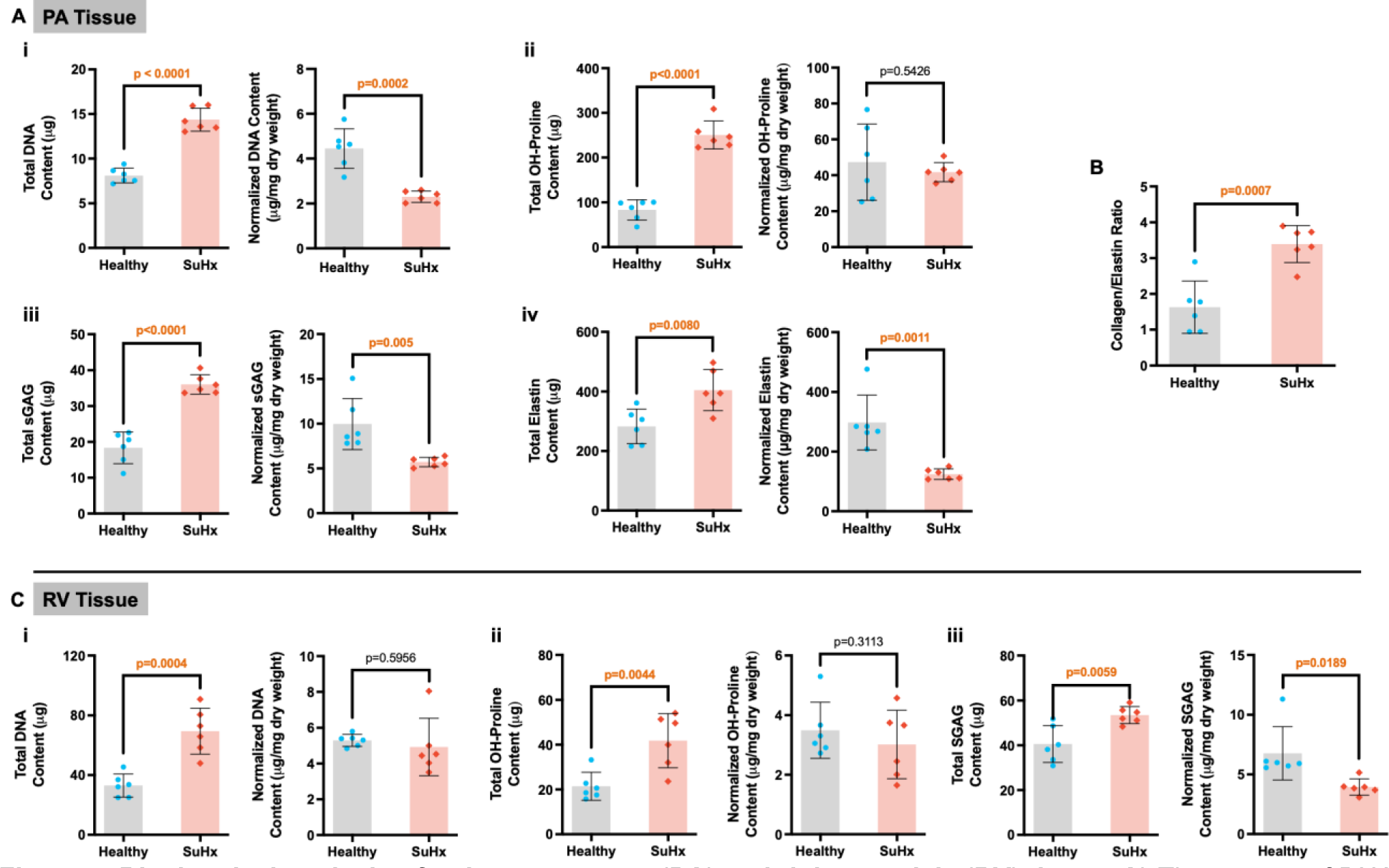
Biochemical analysis of pulmonary artery (PA) and right ventricle (RV) tissue. **A)** The content of DNA (i), hydroxyproline (OH-proline) (ii), sulfated glycosaminoglycan (sGAG) (iii), and elastin (iv) in healthy and sugen hypoxia (SuHx) PA tissues represented as total content and normalized to the tissue dry weight. **B)** The ratio of collagen/elastin contents of PA tissue (OH-proline was considered as 10% of collagen content). **C)** The DNA (i), OH-proline (ii), and sGAG (iii) content in the healthy and SuHx RV tissue, represented in total and normalized to tissue dry weight. Data are presented as mean ± SD with n=6 healthy and n=6 SuHx rats.

### mPA and RV mechanical remodelling

The force-displacement behaviour of mPA and RV tissues is reported as membrane tension-strain instead of stress-strain in the main text and figures of this manuscript. As opposed to stress (force divided by tissue cross-sectional area), membrane tension (force divided by the tissue width) reflects the force carried by the tissue with its full thickness. As both mPA and RV tissues showed significant thickening in PAH, mechanical properties derived based on membrane tension fully reflect how these tissues behave under hemodynamic loading in health and disease, regardless of their thickness, and are more relevant for studying the correlation between tissue mechanical properties and hemodynamic parameters. The stress-strain data are available in the Supplementary Materials (**Figure S1**).

On biaxial mechanical testing, SuHx mPAs showed reduced extensibility and increased high-strain stiffness by 9.9 and 6.7-fold in the longitudinal (p=0.0016) and circumferential (p=0.0005) directions, respectively (**Figure 5A**). In contrast, lower longitudinal and circumferential stiffness were observed in the diseased mPA tissue in the low-strain regime. This phenomenon was observed when load was represented as membrane tension (**Figure 5A-ii**) or stress (**Figure S1B**). More mPA stiffening in the longitudinal direction was associated with a trend for increased mechanical anisotropy (ratio of longitudinal to circumferential stiffness) from 0.9±0.2 in healthy to 1.6±1.1 in SuHx animals. High-strain stiffening of mPA tissue in PAH rats was consistent with tissue fibrosis observed via histological (**Figure 1C-E**) and biochemical (**Figure 4A-ii**) analyses. Reduced PA low-strain stiffness, particularly when calculated based on stress (**Figure S1B**), was associated with lower elastin content normalized to dry weight in SuHx mPA tissue (**Figure 4B-iv**), as elastin mainly governs tissue stiffness in the low-strain regime^24^.

**Figure 5.**
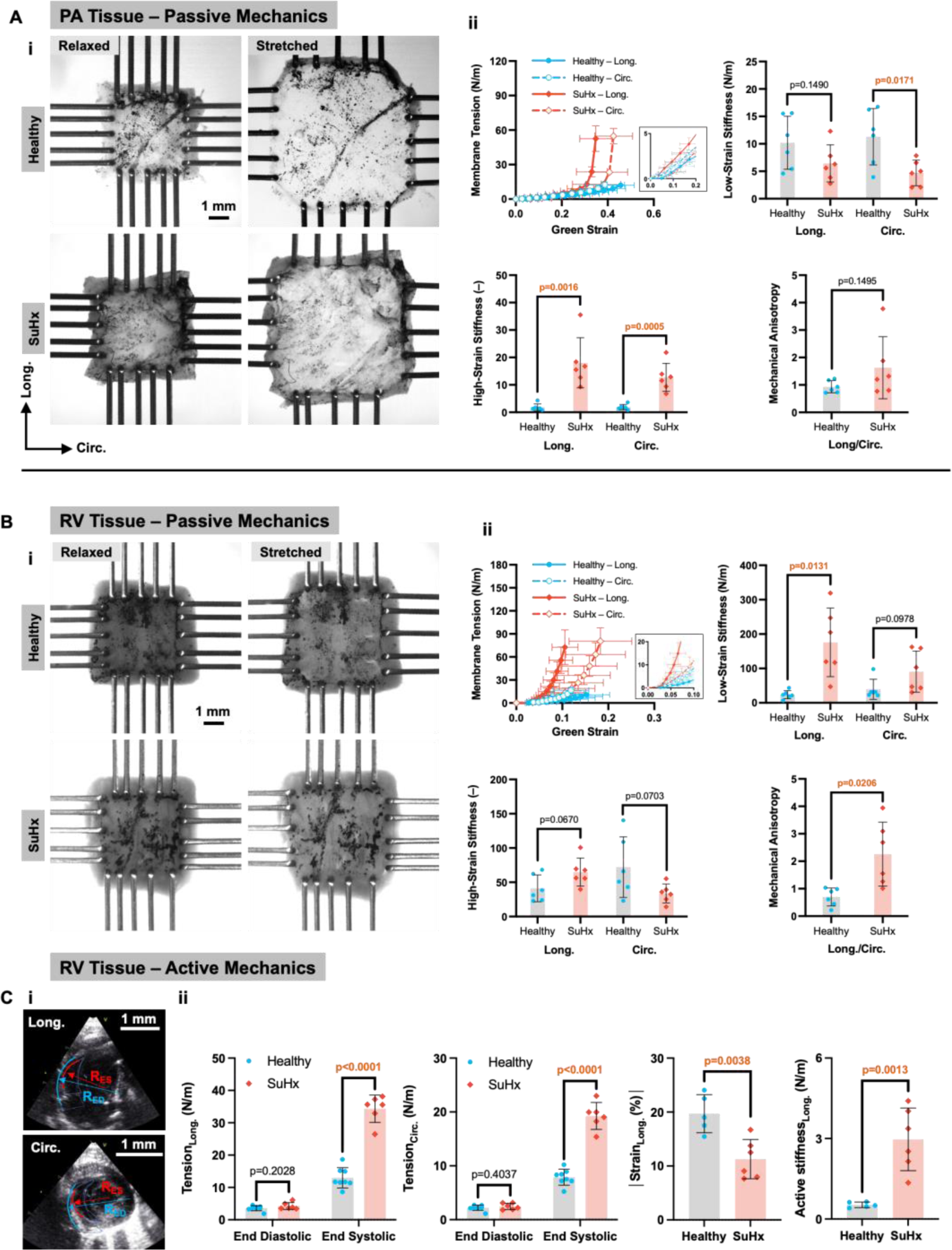
Pulmonary artery (PA) and right ventricle (RV) tissue mechanical properties. **A)** i) Healthy and sugen hypoxia (SuHx) PA tissue in relaxed and stretched states undergoing equibiaxial tensile testing. ii) Membrane tension-strain relationships of healthy and SuHx PA tissues in longitudinal (Long.) and circumferential (Circ.) directions and mechanical characteristics derived from these relationships, including stiffness in the low- and high-strain regions of the tension-strain curve and mechanical anisotropy (Long./Circ. stiffness). As detailed in Supplemental Material, stiffnesses were determined from calculated material constants: C_1_·C_2_ and C_1_·C_3_ to reflect the longitudinal and circumferential low-strain stiffness, respectively [units N/m]; and C_2_ and C_3_ to reflect the longitudinal and circumferential high-strain stiffness, respectively [unitless]. **B)** RV tissues in relaxed and stretched conditions undergoing equibiaxial tensile testing. ii) Passive membrane tension-strain curves in the Long. and Circ. direction of the heart, derived from biaxial mechanical testing of healthy and SuHx RV tissues along with mechanical properties of RV tissues including stiffness in the low- and high-strain regions of the tension-strain curves and mechanical anisotropy. **C)** Echocardiography (i) and cardiac catheterization were used to determine active mechanical behaviour (ii) of RV tissue, including Long. and Circ. tension, Long. strain, and Long. elastic modulus in healthy and SuHx animals in the cardiac cycle. Data are presented as mean ± SD with n=6 healthy and n=6 SuHx rats.

RV free-wall tissue showed increased stiffness in the low-strain regime, significantly in the longitudinal direction from 23.9 ± 12.0 N/m in healthy to 175.9 ± 99.6 N/m in SuHx animals (p=0.013; **Figure 5B**). RV mechanical anisotropy increased from 0.7 ± 0.3 in healthy to 2.3 ± 1.2 in SuHx rats (p=0.021). Higher RV stiffening in the longitudinal direction, and therefore increased tissue anisotropy, reflected the organization of muscle fibres predominantly toward the longitudinal direction, as evidenced in polarized light micrographs (**Figures 3B and C**). This tissue mechanical transformation was observed when load was represented as membrane tension (**Figure 5B-ii**) and stress (**Figure S1C**). RV stiffening was also observed in the high-strain regime in the longitudinal direction from 41.2 ± 19.6 in healthy to 65.0 ± 20.5 in SuHx (p=0.067), which can be explained by the organization of collagen fibres in the longitudinal direction of diseased RV tissues (**Figures 2C-E**).

### RV active mechanical properties

Determined by conductance catheter-pressures and echocardiography (**Figure 5C-i**), end-diastolic active tensions in longitudinal and circumferential directions were similar in healthy and SuHx rats. (**Figures 5C-ii**). Although RVEDP was higher in SuHx rats, their diastolic RV curvature was also higher compared to healthy rats; therefore, the RVEDP/ RV curvature ratio, which governs active tension, remained constant from healthy to SuHx. In contrast, end-systolic tension was significantly higher in SuHx animals by 2.6 and 2.4-fold (p<0.0001) in the longitudinal and circumferential directions, respectively (**Figure 5C-ii**). SuHx rats showed reduced longitudinal strain magnitude by 0.57-fold (p=0.0038) and increased active stiffness by 5.6-fold (p=0.0013) (**Figure 5C-ii**).

### Correlations between PA and RV mechanical properties

Multivariate analysis showed correlations between PA and RV passive mechanical properties, particularly in the longitudinal direction, consistent with mechanical RV-PA coupling (**Figures 6A, 6B, and S3**). PA high-strain longitudinal stiffness correlated with RV low-strain longitudinal stiffness (r=0.82 p=0.0010), RV high-strain longitudinal stiffness (r=0.69; p=0.014), and RV mechanical anisotropy (r=0.69; p=0.013). Additionally, PA mechanical anisotropy correlated with RV low- and high-strain longitudinal stiffness and anisotropy (r=0.60; p=0.039). Correlations were also observed between RV active mechanics and PA and RV passive mechanical properties (**Figures 6C and S4**), particularly between PA high-strain stiffness and RV end-systolic tension in the longitudinal (r=0.86; p=0.0003) and circumferential (r=0.88; p=0.0002) directions (**Figure 6D**). This indicates that high tension levels in the RV myocardium were strongly associated with PA stiffening. Additionally, RV myocardium experiencing higher end-systolic circumferential and longitudinal tension exhibited higher passive mechanical anisotropy (**Figures 6D and S4**).

**Figure 6.**
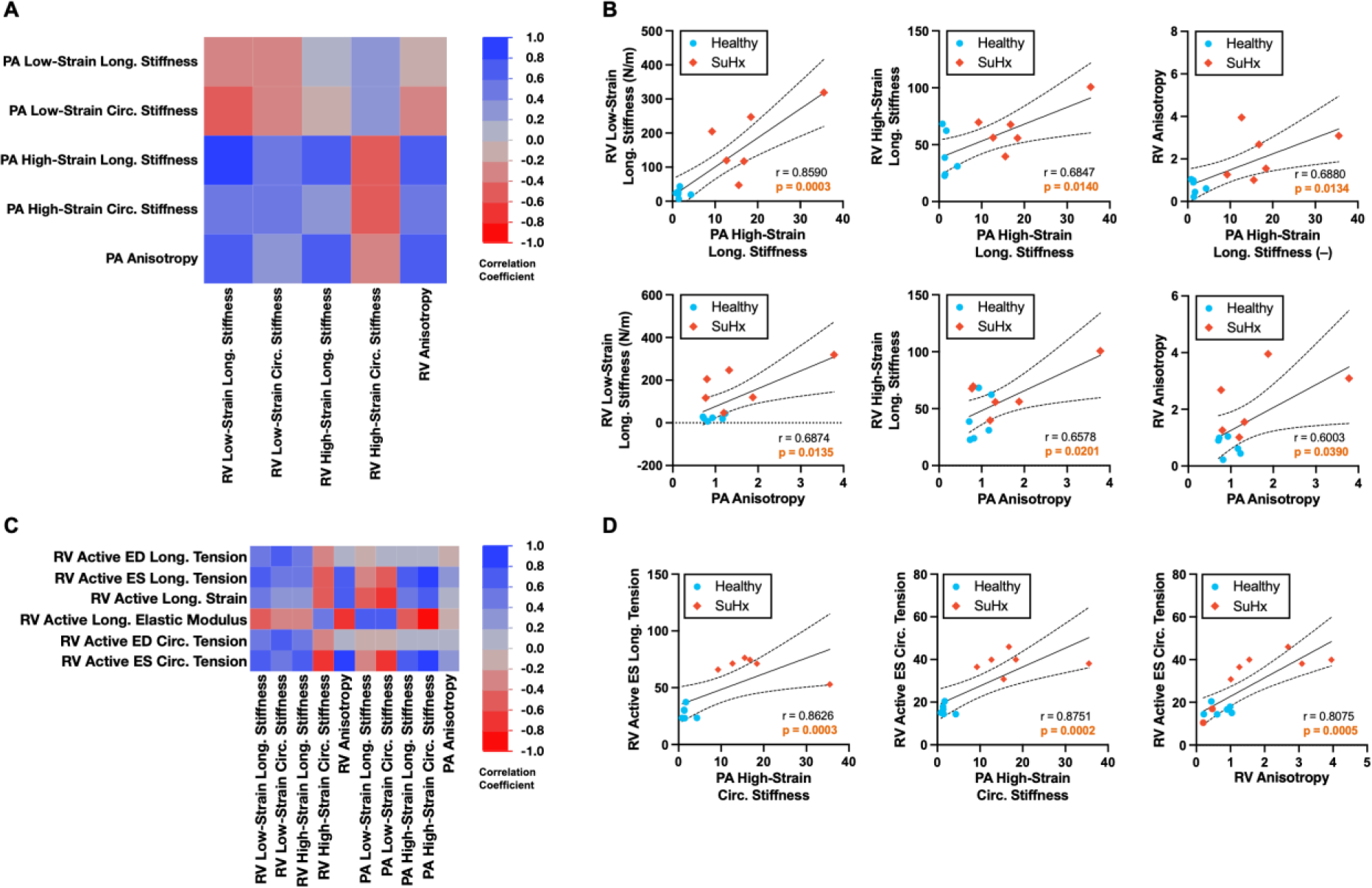
Correlation between pulmonary artery (PA) and right ventricle (RV) tissue mechanics. **A)** Colour map derived from multivariate analysis showing pair-wise correlation of PA and RV passive mechanical properties. **B)** Selected relationships between PA and RV passive mechanical properties. **C)** Colour map representing the correlation between RV active mechanics and RV and PA passive mechanics. **D)** Relationships between RV active tension and PA stiffness as well as RV anisotropy were determined to have the highest correlation coefficients. n=6 healthy and n=6 sugen hypoxia (SuHx) rats.

### Correlation between mPA mechanical remodelling and RV hemodynamics

Multivariate analysis showed correlations between mPA mechanical properties and hemodynamic parameters (**Figures 7A and S5**). In general, these correlations were more pronounced between mPA mechanical properties in the high-strain region, especially in the circumferential direction, and RV hemodynamic function. Functional volume parameters such as SV and CO inversely correlated with PA longitudinal and circumferential stiffness in the high-strain region of the tension-strain curve (**Figure 7B**). In contrast, pressure-related systolic parameters including SW, PRSW and maximum dP/dt directly correlated with mPA high-strain stiffness in the longitudinal and circumferential directions (**Figure 7B**). These mechanical parameters also correlated with RV contractility as reflected by Ees. mPA mechanical properties also correlated with diastolic parameters; notably, minimum dP/dt and the time constant of RV relaxation (tau) inversely correlated with mPA longitudinal and circumferential stiffness in the high-strain region of the tension-strain curve (**Figure 7C**). The PA-RV coupling ratio, Ea/Ees, strongly correlated with mPA low-strain circumferential stiffness (r=-0.93; p=0.0067) and high-strain longitudinal stiffness (r=0.94; p=0.0050) in SuHx rats (**Figure 7D**). RVSP/LVSP and mPA pulse-pressure positively correlated with mPA high-strain stiffness, showing association between the pressure generated by RV (and exerted on the mPA) and mPA stiffening (**Figures 7E, and S5**). In contrast, negative correlations were observed between PA pulsatility and PA high strain stiffness (**Figures 7E and S5**).

**Figure 7.**
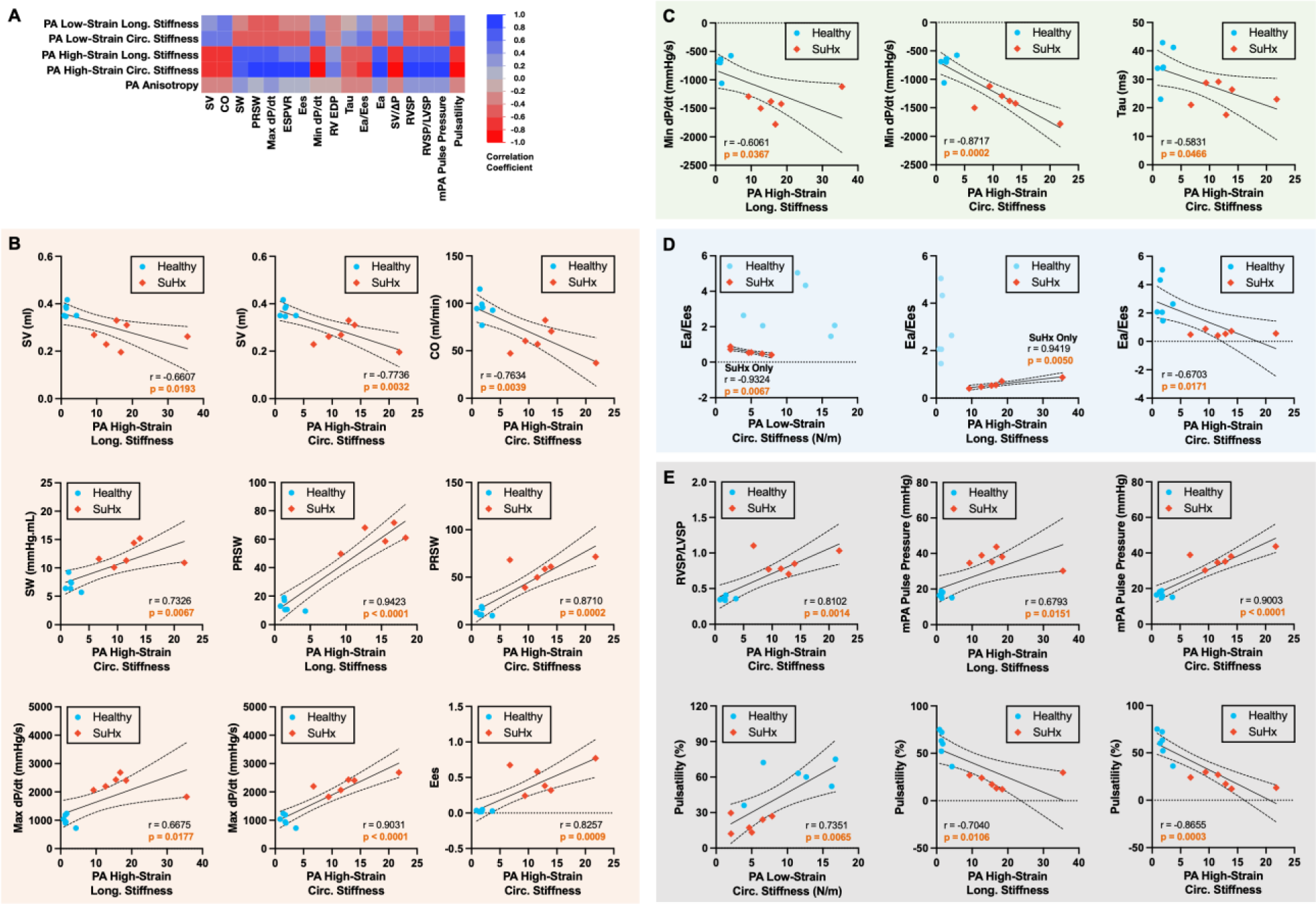
Correlation between pulmonary artery (PA) tissue mechanics and hemodynamic parameters. **A)** Colour map showing the correlation between PA tissue mechanical properties – including stiffness in the longitudinal (Long.) and circumferential (Circ.) directions and mechanical anisotropy – and hemodynamic parameters. **B)** Correlation between PA tissue mechanical properties and systolic hemodynamic parameters, including stroke volume (SV), cardiac output (CO), stroke work (SW), preload recruitable SW (PRSW), max dP/dt, and end-systolic elastance (Ees). **C)** Correlation between PA tissue mechanical properties and diastolic hemodynamic parameters, including min dP/dt and isovolumic relaxation time constant, tau. **D)** Correlation between PA mechanical properties and the PA-RV coupling parameter, arterial elastance (Ea)/Ees. **E)** Correlation between PA mechanical properties and hemodynamic parameters, including right ventricular systolic pressure (RVSP)/ left ventricular systolic pressure (LVSP), main PA (mPA) pulse pressure, and pulsatility. n=6 healthy and n=6 sugen hypoxia (SuHx) rats.

### Correlation between RV passive mechanical remodelling and hemodynamics

RV hemodynamic parameters correlated with RV passive mechanical properties (**Figure 8A**). Volumetric hemodynamic parameters including SV and CO correlated directly with circumferential stiffness but inversely with longitudinal stiffness (**Figures 8A and B**). Therefore, RV mechanical anisotropy (longitudinal/circumferential stiffness) inversely correlated with SV (r=-0.84; p=0.0002) and CO (r=-0.75; p=0.0021) (**Figure 8B**). RV mechanical anisotropy also correlated inversely with diastolic hemodynamic parameters, min dP/dt (r=-0.71; p=0.0046) and tau (r=-0.58; p= 0.031; **Figure 8C**). RVEDP correlated directly with RV stiffness in the low-strain region in the longitudinal (r=0.66; p=0.010) and circumferential (r=0.86; p=0.028) directions (**Figure 8C**). A correlation was also observed between RV anisotropy and the RV-PA coupling ratio, Ea/Ees (r=-0.54; p=0.047; **Figure 8D**). Other hemodynamic parameters, such as RVSP/LVSP and mPA pulse pressure, also correlated with RV mechanics, more significantly with RV anisotropy (**Figure 8E**).

**Figure 8.**
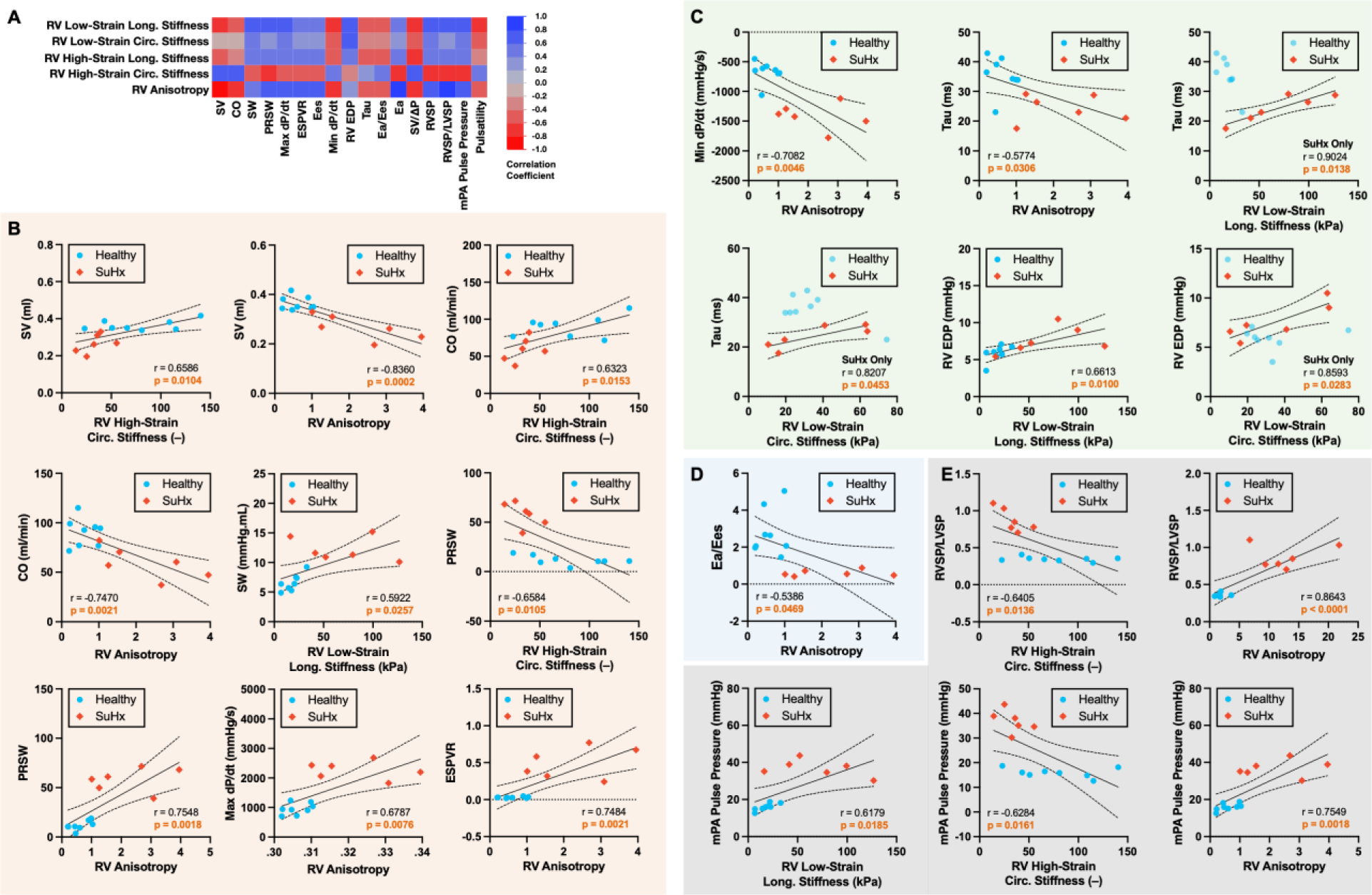
Correlation between right ventricle (RV) tissue passive mechanics and hemodynamic parameters. **A)** Colour map showing the correlation between RV tissue mechanical properties – including mechanical anisotropy and stiffness in the longitudinal (Long.) and circumferential (Circ.) directions – and hemodynamic parameters. **B)** Correlation between RV tissue mechanical properties and systolic hemodynamic parameters, including stroke volume (SV), cardiac output (CO), stroke work (SW), preload recruitable SW (PRSW), max dP/dt, and end-systolic elastance (Ees). **C)** Correlation between RV tissue mechanical properties and diastolic hemodynamic parameters, including min dP/dt, RV end-diastolic pressure (RV EDP), and isovolumic relaxation time constant, tau. **D)** Correlation between RV anisotropy and PA-RV coupling parameter, arterial elastance (Ea)/end-systolic elastance (Ees). **E)** Correlation between RV mechanics and hemodynamic parameters, including right ventricular systolic pressure (RVSP)/ left ventricular systolic pressure (LVSP) and main PA (mPA) pulse pressure. n=6 healthy and n=6 sugen hypoxia (SuHx) rats.

## Discussion

RV-PA coupling is increasingly recognized as an important determinant of RV adaptation to its load, function, and clinical outcomes in PAH^16^. However, the characteristics of the proximal pulmonary conductance vessels and their impact on RV structure, mechanics and function in PAH are incompletely delineated. Consequently, current therapies remain almost entirely focused on the distal pulmonary vasculature, while proximal pulmonary vasculature characteristics are not routinely used for prognostication or as a therapeutic target. The main results of this study show that in PAH: 1) The mPA undergoes significant changes in structure, supported by histology image analysis. These correspond with marked biochemical sGAG deposition and fibrosis and a decrease in the ratio of elastin to collagen; 2) These histological and structural changes are associated with mPA dilatation, thickening and markedly increased stiffness in the longitudinal and circumferential directions; 3) mPA stiffness and remodeling are related to the hemodynamic load, RV-PA coupling and RV mechanics, including RV stiffness and active tension; 4) Changes in RV stiffness and active tension showed a strong association with changes in RV microarchitecture including myofiber orientation, a shift to a trilayer architecture and increased collagen deposition; 5) These changes are associated with RV dysfunction and impaired RV-PA coupling. Taken together, our data suggest that loss of the mPA inherent elastic properties is an important contributor to adverse RV remodeling and RV-PA uncoupling in PAH.

### PA remodeling in PAH

In PAH, most emphasis has naturally been placed on the pathophysiology of the distal pulmonary vasculature. These distal changes lead to increased PVR, determined by the non-pulsatile component of pulmonary blood flow. As evidenced in the current study, distal pulmonary vascular remodeling results in substantial histological changes, remodeling and stiffening of the proximal capacitance vessels. This proximal vascular remodeling contributes to the total vascular impedance faced by the RV, increases RV workload, and consequently metabolism, and ultimately impacts clinical outcomes^15^. Therefore, understanding the structural changes in the tissue (characterized by histological analysis) leading to increased mPA stiffness may inform new therapeutic approaches. Our results show that the mPA undergoes significant dilatation, and wall thickening, with increased sGAG and adventitial collagen deposition and reduced medial elastin. These results are consistent with previous reports in hypoxia-induced PAH models and in patients with PAH, and associated with increased passive PA stiffness^25–29^. However, those reports did not assess the impact on the RV and on RV-PA coupling, which we show are significant^26^.

Elastin and collagen comprise ∼40% of the mPA dry weight in young adults^26,30^, and their quantity and microarchitecture govern the mechanical behaviour of mPA tissue at low and high strains, respectively^31^. Our biochemical analyses showed that mPAs of SuHx rats have an increased collagen-to-elastin ratio as a consequence of collagen production far exceeding elastin production. Additionally, in polarized light micrographs, collagen appeared more organized in fibrils in both circumferential and longitudinal directions in PAH compared to controls. These changes in the content and organization of collagen explain the substantial stiffening of the mPA and its occurrence at lower strains (locus of the ‘heel’ region), in PAH compared to healthy controls. The relative reduction in medial elastin (when normalized to tissue dry weight) was reflected in reduced low-strain stiffness, especially when considering stress-strain (**Figure S1B**) instead of tension-strain curves, in both circumferential and radial directions. This PAH-mediated mechanical remodelling was associated with reduced mPA extensibility and compliance, with much lower strains in SuHx compared to healthy rats. These changes are consistent with an impaired Windkessel effect of the mPA. The reduced compliance of mPA in PAH was also evident in smaller changes in mPA diameter from diastole to systole observed in SuHx rats. Our results are consistent with models of adult and neonatal PAH demonstrating a similar shift in proximal PA elastin and collagen composition which were associated with reduced PA compliance^31,32^. As we will further discuss, this reduction in mPA compliance, and therefore the Windkessel effect, is expected to impair RV-PA coupling efficiency^31^.

The changes in mPA mechanics in PAH corresponded with increases in total DNA, sGAG content, and collagen deposition, consistent with fibroblast proliferation^33,34^. However, the impact of increased sGAG content on mPA stiffening is not well studied. sGAG may impact arterial stiffening by a number of mechanisms including regulation of inflammatory responses and collagen turnover^33^. mPA wall thickening and a reduction in DNA content, when normalized to dry weight in our study, also suggests reduced cell density in relation to non-cellular material. These results may be attributable to edema or to vSMC hypertrophy and muscularization of the medial layer^36^. SMC hypertrophy may further contribute to increased mPA stiffness via mPA thickening and increased vascular tone^37^. However, this relationship has not been clearly delineated and needs further study.

### mPA remodeling and association with RV afterload, flow dynamics, tension, and RV-PA coupling

The RV afterload includes static and oscillatory components, determined by pulmonary vascular resistance, compliance, and wave reflections^38^. The resultant RV wall stress impacts RV work and metabolism^15^. Reduced mPA compliance increases pulse-wave velocities and premature wave reflections, resulting in increased PA systolic pressures and RV pulsatile afterload^39^. This pulsatile component accounts for ∼30% of the total hydraulic load, in which the mPA and proximal branches are primary contributors^12^. Hence, the increased mPA stiffness observed in our PAH rats ‘is anticipated to increase RV impedance, workload and energetics.

Optimal RV-PA coupling refers to the efficient transfer of energy between the RV and the pulmonary vasculature, where work efficiency is optimized. RV-PA coupling is a key determinant of RV adaptation to pressure-loading, and is often measured by the ratio between invasively measured Ea and Ees^16^. Consequently, the Ea/Ees ratio has been used as a reference parameter to assess RV adaptation^40^. Parameters of RV-PA coupling are reduced in end-stage PAH^41^ and mPA stiffening is independently associated with reduced RV EF, itself a parameter of RV-PA coupling^9^. Our results demonstrate an inverse relationship between RV-PA coupling and mPA circumferential stiffness in the high-strain region of the PA tension-strain curve. This suggests that the elastic properties of the mPA impact the efficiency of RV-PA coupling. The elastic properties of the mPA govern its ability to act as a flow reservoir, expanding passively during RV ejection and recoiling to propel blood into the distal vasculature, thereby promoting pulmonary blood flow, while at the same time dampening PA pressures, and minimizing pressure variations in the distal pulmonary arteries (pulsatile dampening) ^42^. Thus, in addition to determining the pulsatile component of RV afterload, the mPA plays a role in dampening pulsatile ejection. The loss of pulsatile dampening that occurs with mPA stiffening affects the pulmonary vasculature distally, in a feed-forward loop. Similar mPA morphological changes to those observed in the current study in hypoxia-induced PAH models were associated with increased flow velocities^42,43^. The increased flow velocities and pulsatile flow into the distal pulmonary vasculature promotes vascular endothelial proliferation, remodeling, and inflammation^43^, all important components of pulmonary vascular remodeling in PAH. These effects occur early, and in experimental PAH models, disruptions in the distal pulmonary vascular internal elastic lamina preceded distal vSMC hypertrophy and endothelial cell proliferation^44,45^.

These mechanical and hemodynamic changes in the mPA have clinical consequences. The ability to buffer pulsatile blood flow at high pressures not only reduces shear stress in the distal pulmonary circuit but also impacts RV stroke work^46,47^. We previously showed in children with PAH, that RV stroke work is increased in relation to clinical status and outcomes; and that RV stroke work also informs on RV-PA de-coupling and worse clinical outcomes^48,49^. We further found in the same SuHx PAH rat model used in the current study that the discrepancy between RV free-wall and septal myocardial work is related to RV fibrosis, wasted work, and inefficiency in PAH^50^. In the current study, we expand on these mechanistic findings, and find that mPA stiffness in the high-strain regime positively correlates with RV stroke-work and end-systolic RV tension. As collagen determines stiffness in the high-strain regime, this suggests that the observed increased mPA fibrosis in our model contributes to reduced mPA expansion during systole and may explain the observed negative correlation between mPA high-strain stiffness and pulsatility^31^. This in turn is expected to increase RV work and energy expenditure, and as detailed above, impact the Windkessel effect^47^. mPA compliance can contribute up to 18-fold more than PVR to RV stroke work^9^. Alterations in the mPA’s elastic properties occur before changes in RV function, and may be an independent contributor to RV dysfunction in PAH^5,9^. In the current study, we show that RV active end-systolic tension was predominantly increased in the longitudinal direction and that PA stiffness in the high-strain regime correlated with load-independent parameters of RV contractility such as Ees and PRSW. Our current results also show that PA stiffness is associated with altered RV wall stress. We and others have previously shown that increased RV wall stress is associated with adverse RV remodeling and dysfunction in RV pressure-loading^51^. Taken together, our multivariate analysis of mPA stiffness in relation to RV hemodynamics show that in PAH, impaired mPA elastic properties increase RV afterload, impacting RV wall stress, tension, micro-structural remodeling and function.

Some statistically significant correlations between PA and RV mechanics or between these and RV function arose from the differences between PAH rats and controls, rather than correlations between parameters within each group. Nonetheless, within group correlations were stronger in the PAH rats, and the results show that the PAH PA and RV operate in vastly different domains from healthy controls. Correlations between PA mechanics and RV-PA coupling were noteworthy in two aspects: 1) The correlation between PA stiffness and the Ea/Ees ratio was tight in PAH rats, but non-existent in healthy controls; and 2) The direction of correlation was opposite in the low versus high-strain regimes. The tight correlation emphasizes the importance of the mPA characteristics on RV-PA coupling - a key determinant of RV adaptation - and suggests that the degree of stiffening impacts RV-PA coupling versus de-coupling.

### RV stiffness and its association with RV remodeling and function

Progressive RV hypertrophy, fibrosis, and dilatation contribute to impaired RV-PA coupling and development of RV failure^6,38,52^. The increased RV workload further triggers aberrant RV metabolism which drives RV failure^53^. The low and high strain regions of the RV-tension strain curve represent regions where myofibers and collagen fibers, respectively, determine RV myocardial stiffness^54^. Our results show that SuHx-induced PAH led to increased RV low-strain longitudinal stiffness. This suggests important changes in the longitudinal vector of RV mechanics which could be attributed to more longitudinally aligned RV myofibers in PAH rats. These results are consistent with previous work that have shown increased longitudinal stiffness in the low-strain regime of the RV stress-strain curve^55^. These changes have functional consequences in that in our study, increased RV myofiber anisotropy was associated with reduced RV longitudinal strain, SV, and RV-PA coupling. This emphasizes the importance of longitudinal myofiber stiffness to RV dysfunction. The severity of the increased load also seems important, as in more adaptive RV pressure-loading models, RV myofiber arrangement, assessed by magnetic resonance diffusion tensor imaging, was not different compared to healthy controls^56^. Similar results have been found by other studies in the PAB model, although, using computational modeling, those studies found increased microstructural disorganization (dispersion) in association with increased RV wall thickness, stiffness, pro-fibrotic gene expression and collagen deposition^57^. These findings align well with our own prior work in the PAB model where we found RV hypertrophy, and robust pro-fibrotic signaling and fibrosis^22,58^. In those studies, increased RV free-wall stiffness was associated with increased collagen deposition and contributed to increased RV chamber stiffness as reflected by pressure-volume loop analysis^57^.

Our current results showing increased RV myofiber anisotropy and increased longitudinal stiffness add to the aforementioned clinical implications in that commonly used, and prognostically important, parameters of longitudinal RV function such as tricuspid annular plane systolic excursion (TAPSE), tissue Doppler velocities, and longitudinal strain are reduced in PAH^59^. To maintain RV-PA coupling, the RV compensates for the reduced longitudinal function by adapting a more radial/circumferential contraction^60,61^. Our results suggest that this may occur because of a change in myocyte orientation and/or anisotropy. Indeed, we observed development of a tri-layer architecture in the PAH RV, reminiscent of a normal LV, which has a dominant circumferential contraction pattern to allow ventricular-arterial coupling in the high-pressured systemic circulation. Our empirical observations support results from computational modeling studies, where in response to PAH, the earliest adaptations were increased myofiber contractility, and an increase in RV passive stiffness, as we observed in our study^62^. These occur to maintain cardiac output and RV-PA coupling^62^. However, in that computational study, reorientation of myofibers toward the longitudinal (apex-to-base) direction resulted in impaired RV free-wall contractile patterns, ultimately resulting in a maladaptive response^61^. Thus, despite increased intrinsic myofiber contractility, the changes we observed in RV architecture and fiber orientation can result in reduced effective RV function and EF^62^.

RV stiffening affects both systolic and diastolic function. Increased RV diastolic stiffness in patients with PAH results from increased RV cardiomyocyte sarcomere stiffening and from increased collagen deposition^63^. We observed correlations between RV stiffness in the low-strain regime with tau and with RVEDP, suggesting that both active relaxation in early diastole and passive chamber stiffness at end-diastole are affected. We found that active RV end-diastolic wall tension was low and comparable to controls. This may explain the correlation between passive RV stiffness in the low-strain regime and early and late diastolic parameters (tau and RVEDP, respectively). Nonetheless, passive RV stiffness was significantly increased in the low strain regime, especially in the longitudinal direction. Taken together, these results may support the notion that intrinsic myofiber stiffness, more than fibrosis, may be a key determinant of diastolic impairment in PAH^63^, as the low-strain regime is thought to be determined by the myocyte compartment whereas the high-strain regime is predominantly determined by collagen^64,65^. This result is also in keeping with the high RV mechanical anisotropy in the PAH rats, which reflects myofiber arrangement. Indeed, the active RV elastic modulus was markedly elevated in PAH rats in the longitudinal direction. Overall, our results are in keeping with Rain et al. who found that the myocyte compartment determined stiffness at lower RV pressures, with fibrosis playing an increasingly important role in determining active stiffness with greater disease severity and higher RV pressures^64^. The increased PA and RV stiffness are clinically relevant as end-diastolic elastance is a predictor of response to therapy in patients with PAH^40^. The increases in RV pressures and stiffness, also affect right atrial (RA) stiffness, because of increased RA cardiomyocyte stiffness and increased interstitial fibrosis^66^. This is relevant as echocardiographic studies found that in patients with suspected PAH, the peak lateral tricuspid annulus systolic velocity/right atrial area index ratio (S’/RAAi) predicted elevated RV end-diastolic elastance and survival^67^.

### Clinical implications

The correlation of mPA stiffness with RV stiffness has major clinical implications as discussed above. In particular, the strong correlation between mPA stiffness and RV-PA coupling, as determined by the Ea/Ees ratio would imply correlation with clinical outcomes. Thus, although development of mPA stiffening results from the distal vascular remodeling, our results suggest that the mPA stiffening itself significantly impacts RV function and hence, by extension, clinical outcomes. Therefore, our pre-clinical results suggest that mPA compliance should be assessed clinically. Indeed, we, and others, have shown that RV capacitance is related to outcomes in patients with PAH^68^. Moreover, PA capacitance and/ or mPA or branch PA compliance, derivates of pulsatility and related to mPA stiffness, as we demonstrate in the current study, can be assessed non-invasively by echocardiography or MRI, thereby making their measurement clinically feasible and perhaps more accessible than measurement of PVR^69^. Moreover, with the evolution of ultrafast ultrasound (UFUS), myocardial and mPA stiffness can be assessed non-invasively^70^. Further advances in UFUS may also inform on RV fiber orientation in vivo, which may inform on RV adaptation^71^. From a therapeutic standpoint, treatment of the distal-intra-pulmonary artery disease will remain the cornerstone of PAH therapy and will impact the mPA stiffness. The results form the basis for further studies that may direct the treatment of mPA stiffness, for example by restoring the elastin/collagen ratio, or reducing excess sGAG deposition. Our results also suggest that therapies to address RV fibrosis or to modulate its stiffness, independent of the distal pulmonary vascular remodeling would be valuable.

### Limitations

The SuHx model utilizes SU5416, a VEGFR2 inhibitor, coupled with hypoxia to induce PAH. VEGFR2 mediates angiogenesis in the heart, which may affect RV remodeling^18^. Because biochemical tissue assays cannot be done on the same tissues where stiffness is assessed, histology and biochemical assays were done on additional animals. We did not repeat echocardiography and catheterization in those animals, as these were performed in animals whose tissues were used for stiffness assessment. Approximation of RV active tension was calculated by Laplace’s law using echocardiography measurements and assuming a constant and spherical RV curvature. Inaccuracies in endocardial traces can affect measures of the radius of curvature and could have affected results. Although these assumptions may be overly simplistic^72^, we previously reported excellent agreement between computer-modelled wall tension and wall tension calculated by echocardiography-derived measurements^51^. For biaxial mechanical testing, we assumed uniform stress and strain fields in the tissue samples during stretch cycles. This assumption undermines the heterogeneity of test samples, and the method derives lumped mechanical properties that might not reflect the mechanics of focal regions of mPA and RV tissues. We utilized a rat model in this study. Tian and colleagues found that proximal PA remodeling in calves is different from that in rats, so that species-related effects may be important and needs to be considered when translating results to humans^29^. Lastly, we studied a single time-point at 6-weeks. Further studies are required to evaluate temporal changes in PA and RV stiffening, and how these relate to progressive RV dysfunction and RV-PA decoupling.

### Conclusions

Reduced mPA compliance and impaired RV-PA coupling are increasingly recognized as risk factors for adverse clinical outcomes in PAH^6^. Our data suggest that distal pulmonary vascular remodeling induces geometrical and structural changes in the mPA that increase its stiffness. The mPA disease in turn is associated with adverse RV remodeling and changes in its micro-architecture associated with increased RV wall stiffness, stress, tension, and dysfunction, particularly in the longitudinal direction. Together, these are associated with impaired RV-PA coupling. Therapies to reduce mPA and/or RV myocardial stiffness and myofiber remodeling may prevent or slow adverse RV remodeling and improve RV function in PAH thereby improving morbidity and mortality in this severe condition.

## List of Abbreviations

CO: Cardiac output
CO: Cardiac ouput
Ea: Arterial elastance
EDP: End-diastolic pressure
EDPVR: End-disastolic pressure volume relations
Ees: End-systolic elastance
LV: Left ventricle
LVSP: Left ventricular systolic pressure
mPA: Main pulmonary artery
PAH: Pulmonary arterial hypertension
PRSW: Preload recruitable stroke work
PSR: Picrosirius red
PVR: Pulmonary vascular resistance
RV: Right ventricle
RV-PA: Right ventricular-pulmonary arterial
RVSP: Right ventricular systolic pressure
sGAGs: Sulfated glycosaminoglycans
SuHx: Sugen + hypoxia
SV: Stroke volume
SW: Stroke work
vSMC: Vascular smooth muscle cell

## Acknowledgements

This work was supported by the Canadian Institute of Health Research (PHT-169037), the Translational Biology and Engineering Program of the Ted Rogers Centre for Heart Research, and an NSERC Canada Graduate Scholarship-Doctoral and an Ontario Graduate Scholarship to BM.

## References

1 Seeger, W. et al. Pulmonary hypertension in chronic lung diseases. J Am Coll Cardiol 62, D109–116 (2013). 10.1016/j.jacc.2013.10.036

2 Maron, B. et al. Pulmonary vascular resistance and clinical outcomes in patients with pulmonary hypertension: a retrospective cohort study. Lancet Respir Med 8, 873–884 (2020). 10.1016/S2213-2600(20)30317-9

3 Gan, C. et al. Noninvasively assessed pulmonary artery stiffness predicts mortality in pulmonary arterial hypertension. Chest 132, 1906–1912 (2007). 10.1378/chest.07-1246

4 Swift, A. et al. Pulmonary artery relative area change detects mild elevations in pulmonary vascular resistance and predicts adverse outcome in pulmonary hypertension. Invest Radiol 47, 571–577 (2012). 10.1097/RLI.0b013e31826c4341

5 Sanz, J. et al. Evaluation of pulmonary artery stiffness in pulmonary hypertension with cardiac magnetic resonance. JACC Cardiovasc Imaging 2, 286–295 (2009). 10.1016/j.jcmg.2008.08.007

6 Mahapatra, S., Nishimura, R., Sorajja, P., Cha, S. & McGoon, M. Relationship of pulmonary arterial capacitance and mortality in idiopathic pulmonary arterial hypertension. J Am Coll Cardiol 47, 799–803 (2006). 10.1016/j.jacc.2005.09.054

7 Al-Naamani, N., Preston, I., Paulus, J., Hill, N. & Roberts, K. Pulmonary Arterial Capacitance Is an Important Predictor of Mortality in Heart Failure With a Preserved Ejection Fraction. JACC Heart Fail 3, 467–474 (2015). 10.1016/j.jchf.2015.01.013

8 Malhotra, R. et al. Pulmonary Vascular Distensibility Predicts Pulmonary Hypertension Severity, Exercise Capacity, and Survival in Heart Failure. Circ Heart Fail 9 (2016). 10.1161/CIRCHEARTFAILURE.115.003011

9 Stevens, G. et al. RV dysfunction in pulmonary hypertension is independently related to pulmonary artery stiffness. JACC Cardiovasc Imaging 5, 378–387 (2012). 10.1016/j.jcmg.2011.11.020

10 Campo, A. et al. Hemodynamic predictors of survival in scleroderma-related pulmonary arterial hypertension. Am J Respir Crit Care Med 182, 252–260 (2010). 10.1164/rccm.200912-1820OC

11 Mahapatra, S., Nishimura, R., Oh, J. & McGoon, M. The prognostic value of pulmonary vascular capacitance determined by Doppler echocardiography in patients with pulmonary arterial hypertension. J Am Soc Echocardiogr 19, 1045–1050 (2006). 10.1016/j.echo.2006.03.008

12 Saouti, N., Westerhof, N., Postmus, P. & Vonk-Noordegraaf, A. The arterial load in pulmonary hypertension. Eur Respir Rev 19, 197–203 (2010). 10.1183/09059180.00002210

13 Vélez-Rendón, D., Zhang, X., Gerringer, J. & Valdez-Jasso, D. Compensated right ventricular function of the onset of pulmonary hypertension in a rat model depends on chamber remodeling and contractile augmentation. Pulm Circ 8, 2045894018800439 (2018). 10.1177/2045894018800439

14 Bogaard, H., Abe, K., Vonk Noordegraaf, A. & Voelkel, N. The right ventricle under pressure: cellular and molecular mechanisms of right-heart failure in pulmonary hypertension. Chest 135, 794–804 (2009). 10.1378/chest.08-0492

15 Vonk-Noordegraaf, A. et al. Right heart adaptation to pulmonary arterial hypertension: physiology and pathobiology. J Am Coll Cardiol 62, D22–33 (2013). 10.1016/j.jacc.2013.10.027

16 Vanderpool, R. et al. RV-pulmonary arterial coupling predicts outcome in patients referred for pulmonary hypertension. Heart 101, 37–43 (2015). 10.1136/heartjnl-2014-306142

17 Lahm, T. et al. Assessment of Right Ventricular Function in the Research Setting: Knowledge Gaps and Pathways Forward. An Official American Thoracic Society Research Statement. Am J Respir Crit Care Med 198, e15–e43 (2018). 10.1164/rccm.201806-1160ST

18 Vitali, S. et al. The Sugen 5416/hypoxia mouse model of pulmonary hypertension revisited: long-term follow-up. Pulm Circ 4, 619–629 (2014). 10.1086/678508

19 Franco, C. et al. Discoidin domain receptor 1 (ddr1) deletion decreases atherosclerosis by accelerating matrix accumulation and reducing inflammation in low-density lipoprotein receptor-deficient mice. Circ Res 102, 1202–1211 (2008). 10.1161/CIRCRESAHA.107.170662

20 Labrosse, M., Jafar, R., Ngu, J. & Boodhwani, M. Planar biaxial testing of heart valve cusp replacement biomaterials: Experiments, theory and material constants. Acta Biomater 45, 303–320 (2016). 10.1016/j.actbio.2016.08.036

21 Sacks, M. A method for planar biaxial mechanical testing that includes in-plane shear. J Biomech Eng 121, 551–555 (1999). 10.1115/1.2835086

22 Akazawa, Y. et al. Pulmonary artery banding is a relevant model to study the right ventricular remodeling and dysfunction that occurs in pulmonary arterial hypertension. J Appl Physiol (1985) 129, 238–246 (2020). 10.1152/japplphysiol.00148.2020

23 Pacher, P., Nagayama, T., Mukhopadhyay, P., Bátkai, S. & Kass, D. Measurement of cardiac function using pressure-volume conductance catheter technique in mice and rats. Nat Protoc 3, 1422–1434 (2008). 10.1038/nprot.2008.138

24 Carta, L. et al. Discrete contributions of elastic fiber components to arterial development and mechanical compliance. Arterioscler Thromb Vasc Biol 29, 2083–2089 (2009). 10.1161/ATVBAHA.109.193227

25 Drexler, E. et al. Stiffening of the Extrapulmonary Arteries From Rats in Chronic Hypoxic Pulmonary Hypertension. J Res Natl Inst Stand Technol 113, 239–249 (2008). 10.6028/jres.113.018

26 Wang, Z. & Chesler, N. Role of collagen content and cross-linking in large pulmonary arterial stiffening after chronic hypoxia. Biomech Model Mechanobiol 11, 279–289 (2012). 10.1007/s10237-011-0309-z

27 Hoffmann, J. et al. Compartment-specific expression of collagens and their processing enzymes in intrapulmonary arteries of IPAH patients. Am J Physiol Lung Cell Mol Physiol 308, L1002–1013 (2015). 10.1152/ajplung.00383.2014

28 Poiani, G. et al. Collagen and elastin metabolism in hypertensive pulmonary arteries of rats. Circ Res 66, 968–978 (1990). 10.1161/01.res.66.4.968

29 Tian, L. et al. Linked opening angle and histological and mechanical aspects of the proximal pulmonary arteries of healthy and pulmonary hypertensive rats and calves. Am J Physiol Heart Circ Physiol 301, H1810–1818 (2011). 10.1152/ajpheart.00025.2011

30 Mackay, E., Banks, J., Sykes, B. & Lee, G. Structural basis for the changing physical properties of human pulmonary vessels with age. Thorax 33, 335–344 (1978). 10.1136/thx.33.3.335

31 Lammers, S. et al. Changes in the structure-function relationship of elastin and its impact on the proximal pulmonary arterial mechanics of hypertensive calves. Am J Physiol Heart Circ Physiol 295, H1451–1459 (2008). 10.1152/ajpheart.00127.2008

32 Ooi, C., Wang, Z., Tabima, D., Eickhoff, J. & Chesler, N. The role of collagen in extralobar pulmonary artery stiffening in response to hypoxia-induced pulmonary hypertension. Am J Physiol Heart Circ Physiol 299, H1823–1831 (2010). 10.1152/ajpheart.00493.2009

33 Papakonstantinou, E. & Karakiulakis, G. The ‘sweet’ and ‘bitter’ involvement of glycosaminoglycans in lung diseases: pharmacotherapeutic relevance. Br J Pharmacol 157, 1111–1127 (2009). 10.1111/j.1476-5381.2009.00279.x

34 Panzhinskiy, E., Zawada, W., Stenmark, K. & Das, M. Hypoxia induces unique proliferative response in adventitial fibroblasts by activating PDGFβ receptor-JNK1 signalling. Cardiovasc Res 95, 356–365 (2012). 10.1093/cvr/cvs194

35 Chai, S. et al. Overexpression of hyaluronan in the tunica media promotes the development of atherosclerosis. Circ Res 96, 583–591 (2005). 10.1161/01.RES.0000158963.37132.8b

36 Wilson, J., Yu, J., Taylor, L. & Polgar, P. Hyperplastic Growth of Pulmonary Artery Smooth Muscle Cells from Subjects with Pulmonary Arterial Hypertension Is Activated through JNK and p38 MAPK. PLoS One 10, e0123662 (2015). 10.1371/journal.pone.0123662

37 Armentano, R. et al. Smart smooth muscle spring-dampers. Smooth muscle smart filtering helps to more efficiently protect the arterial wall. IEEE Eng Med Biol Mag 26, 62–70 (2007). 10.1109/memb.2007.289123

38 Wang, Z. & Chesler, N. Pulmonary vascular wall stiffness: An important contributor to the increased right ventricular afterload with pulmonary hypertension. Pulm Circ 1, 212–223 (2011). 10.4103/2045-8932.83453

39 Castelain, V. et al. Pulmonary artery pulse pressure and wave reflection in chronic pulmonary thromboembolism and primary pulmonary hypertension. J Am Coll Cardiol 37, 1085–1092 (2001). 10.1016/s0735-1097(00)01212-2

40 Vanderpool, R. et al. The Right Ventricular-Pulmonary Arterial Coupling and Diastolic Function Response to Therapy in Pulmonary Arterial Hypertension. Chest 161, 1048–1059 (2022). 10.1016/j.chest.2021.09.040

41 Tello, K. et al. Reserve of Right Ventricular-Arterial Coupling in the Setting of Chronic Overload. Circ Heart Fail 12, e005512 (2019). 10.1161/CIRCHEARTFAILURE.118.005512

42 Tan, W., Madhavan, K., Hunter, K., Park, D. & Stenmark, K. Vascular stiffening in pulmonary hypertension: cause or consequence? (2013 Grover Conference series). Pulm Circ 4, 560–580 (2014). 10.1086/677370

43 Li, M., Scott, D., Shandas, R., Stenmark, K. & Tan, W. High pulsatility flow induces adhesion molecule and cytokine mRNA expression in distal pulmonary artery endothelial cells. Ann Biomed Eng 37, 1082–1092 (2009). 10.1007/s10439-009-9684-3

44 Todorovich-Hunter, L. et al. Increased pulmonary artery elastolytic activity in adult rats with monocrotaline-induced progressive hypertensive pulmonary vascular disease compared with infant rats with nonprogressive disease. Am Rev Respir Dis 146, 213–223 (1992). 10.1164/ajrccm/146.1.213

45 Maruyama, K. et al. Chronic hypoxic pulmonary hypertension in rats and increased elastolytic activity. Am J Physiol 261, H1716–1726 (1991). 10.1152/ajpheart.1991.261.6.H1716

46 Westerhof, N., Lankhaar, J. & Westerhof, B. The arterial Windkessel. Med Biol Eng Comput 47, 131–141 (2009). 10.1007/s11517-008-0359-2

47 Hunter, K., Lammers, S. & Shandas, R. Pulmonary vascular stiffness: measurement, modeling, and implications in normal and hypertensive pulmonary circulations. Compr Physiol 1, 1413–1435 (2011). 10.1002/cphy.c100005

48 Di Maria, M. et al. RV stroke work in children with pulmonary arterial hypertension: estimation based on invasive haemodynamic assessment and correlation with outcomes. Heart 100, 1342–1347 (2014). 10.1136/heartjnl-2013-305298

49 Di Maria, M. et al. Parameters of Right Ventricular Function Reveal Ventricular-Vascular Mismatch as Determined by Right Ventricular Stroke Work versus Pulmonary Vascular Resistance in Children with Pulmonary Hypertension. J Am Soc Echocardiogr 33, 218–225 (2020). 10.1016/j.echo.2019.09.013

50 Ebata, R. et al. Asymmetric Regional Work Contributes to Right Ventricular Fibrosis, Inefficiency, and Dysfunction in Pulmonary Hypertension versus Regurgitation. J Am Soc Echocardiogr 34, 537–550.e533 (2021). 10.1016/j.echo.2020.12.011

51 Gold, J., Akazawa, Y., Sun, M., Hunter, K. & Friedberg, M. Relation between right ventricular wall stress, fibrosis, and function in right ventricular pressure loading. Am J Physiol Heart Circ Physiol 318, H366–H377 (2020). 10.1152/ajpheart.00343.2019

52 Thenappan, T. et al. The Critical Role of Pulmonary Arterial Compliance in Pulmonary Hypertension. Ann Am Thorac Soc 13, 276–284 (2016). 10.1513/AnnalsATS.201509-599FR

53 Archer, S., Fang, Y., Ryan, J. & Piao, L. Metabolism and bioenergetics in the right ventricle and pulmonary vasculature in pulmonary hypertension. Pulm Circ 3, 144–152 (2013). 10.4103/2045-8932.109960

54 Hill, M. et al. Structural and mechanical adaptations of right ventricle free wall myocardium to pressure overload. Ann Biomed Eng 42, 2451–2465 (2014). 10.1007/s10439-014-1096-3

55 Jang, S. et al. Biomechanical and Hemodynamic Measures of Right Ventricular Diastolic Function: Translating Tissue Biomechanics to Clinical Relevance. J Am Heart Assoc 6 (2017). 10.1161/JAHA.117.006084

56 Nielsen, E. et al. Normal right ventricular three-dimensional architecture, as assessed with diffusion tensor magnetic resonance imaging, is preserved during experimentally induced right ventricular hypertrophy. Anat Rec (Hoboken) 292, 640–651 (2009). 10.1002/ar.20873

57 Kakaletsis, S. et al. Untangling the mechanisms of pulmonary arterial hypertension-induced right ventricular stiffening in a large animal model. Acta Biomater 171, 155–165 (2023). 10.1016/j.actbio.2023.09.043

58 Sun, M. et al. Experimental Right Ventricular Hypertension Induces Regional β1-Integrin-Mediated Transduction of Hypertrophic and Profibrotic Right and Left Ventricular Signaling. J Am Heart Assoc 7 (2018). 10.1161/JAHA.117.007928

59 Sachdev, A. et al. Right ventricular strain for prediction of survival in patients with pulmonary arterial hypertension. Chest 139, 1299–1309 (2011). 10.1378/chest.10-2015

60 Pettersen, E. et al. Contraction pattern of the systemic right ventricle shift from longitudinal to circumferential shortening and absent global ventricular torsion. J Am Coll Cardiol 49, 2450–2456 (2007). 10.1016/j.jacc.2007.02.062

61 Mendiola, E. et al. Right Ventricular Architectural Remodeling and Functional Adaptation in Pulmonary Hypertension. Circ Heart Fail 16, e009768 (2023). 10.1161/CIRCHEARTFAILURE.122.009768

62 Avazmohammadi, R. et al. Interactions Between Structural Remodeling and Hypertrophy in the Right Ventricle in Response to Pulmonary Arterial Hypertension. J Biomech Eng 141, 0910161–09101613 (2019). 10.1115/1.4044174

63 Rain, S. et al. Right ventricular diastolic impairment in patients with pulmonary arterial hypertension. Circulation 128, 2016–2025, 2011-2010 (2013). 10.1161/CIRCULATIONAHA.113.001873

64 Rain, S. et al. Right Ventricular Myocardial Stiffness in Experimental Pulmonary Arterial Hypertension: Relative Contribution of Fibrosis and Myofibril Stiffness. Circ Heart Fail 9, e002636 (2016). 10.1161/CIRCHEARTFAILURE.115.002636

65 Avazmohammadi, R., Hill, M., Simon, M., Zhang, W. & Sacks, M. A novel constitutive model for passive right ventricular myocardium: evidence for myofiber-collagen fiber mechanical coupling. Biomech Model Mechanobiol 16, 561–581 (2017). 10.1007/s10237-016-0837-7

66 Wessels, J. et al. Right Atrial Adaptation to Precapillary Pulmonary Hypertension: Pressure-Volume, Cardiomyocyte, and Histological Analysis. J Am Coll Cardiol 82, 704–717 (2023). 10.1016/j.jacc.2023.05.063

67 Yogeswaran, A. et al. Echocardiographic evaluation of right ventricular diastolic function in pulmonary hypertension. ERJ Open Res 9 (2023). 10.1183/23120541.00226-2023

68 Sajan, I. et al. Pulmonary arterial capacitance in children with idiopathic pulmonary arterial hypertension and pulmonary arterial hypertension associated with congenital heart disease: relation to pulmonary vascular resistance, exercise capacity, and survival. Am Heart J 162, 562–568 (2011). 10.1016/j.ahj.2011.06.014

69 Friedberg, M., Feinstein, J. & Rosenthal, D. Noninvasive assessment of pulmonary arterial capacitance by echocardiography. J Am Soc Echocardiogr 20, 186–190 (2007). 10.1016/j.echo.2006.08.009

70 Villemain, O. et al. Ultrafast Ultrasound Imaging in Pediatric and Adult Cardiology: Techniques, Applications, and Perspectives. JACC Cardiovasc Imaging 13, 1771–1791 (2020). 10.1016/j.jcmg.2019.09.019

71 Papadacci, C. et al. Imaging the dynamics of cardiac fiber orientation in vivo using 3D Ultrasound Backscatter Tensor Imaging. Sci Rep 7, 830 (2017). 10.1038/s41598-017-00946-7

72 Wong, A. & Rautaharju, P. Stress distribution within the left ventricular wall approximated as a thick ellipsoidal shell. Am Heart J 75, 649–662 (1968). 10.1016/0002-8703(68)90325-6

