## Supplemental Materials for "Adverse structural and mechanical remodelling of main pulmonary artery in experimental pulmonary arterial hypertension is associated with impaired right ventricle-pulmonary artery coupling and function"

*Equal contribution

***Biochemical analysis:*** The excised tissues were washed with Dulbecco's phosphate-buffered saline (DPBS) +/+ (Corning, USA, cat# 21-030-CV), placed in pre-weighed centrifuge vials, flash-frozen in liquid nitrogen, and freeze-dried for all biochemical assays. For DNA, OH-proline, and sGAG assays, each sample was digested in 250 µL of papain digestion buffer containing 35 mM ammonium acetate (Sigma-Aldrich, USA, cat# A7330), 1 mM ethylenediaminetetraacetic acid (BioShop, Canada, cat# EDT111), 2 mM DL-dithiothreitol (Sigma-Aldrich, USA, cat# D9779), and 80 µg/mL papain (Sigma-Aldrich, USA, cat# P3125) at pH 6.2 on a heat block at 65°C for 72 hours. For OH-proline assay, acid hydrolysis was performed using 6.0 N HCl (Sigma-Aldrich, USA, cat# 2104) at 110°C for 18 hours. Subsequently, neutralization was carried out using 5.7 N NaOH (VWR International, cat# BDH7247-1). If necessary, the samples were diluted with deionized water. 100 µL of the diluted samples and the standard OH-proline stock solutions were sequentially mixed with 0.05 N chloramine-T (Sigma-Aldrich, USA, cat# 857319-), 3.15 N perchloric acid (Sigma-Aldrich, USA, cat# 311421), and Ehrlich's Reagent (Sigma-Aldrich, USA, cat# 03891). The absorbance of the resulting solution was measured at 560 nm. For the sGAG assay, dimethylmethylene blue (Sigma-Aldrich, USA, cat# 341088) solution was added to the diluted digested samples and chondroitin sulphate sodium salt (Sigma-Aldrich, USA, cat# C9819) solutions as standard reference. The sGAG content was determined by their absorbance at 525 nm, compared to those of reference solutions. For the DNA assay, 0.1 µg/mL Hoechst 33258 (Invitrogen, USA, cat# H3569) DNA stain in a buffer containing 10 mM Trizma base (Sigma-Aldrich, USA, cat# T6066), 0.1 mM NaCl, and 1 mM EDTA at pH 7.4 was added to the digested samples. DNA content was calculated against a calf thymus DNA standard. The excitation and emission of samples were measured at 350 and 450 nm, respectively. Insoluble elastin content was determined using Fastin elastin assay (Accurate Chemical & Scientific Corp., USA, cat# CLRFPD). Freeze-dried samples were digested in 0.25 M oxalic acid (BioShop, Canada, cat# OXA7272) at 100°C for one hour. The supernatant was collected, the oxalic acid treatment was repeated two more times to ensure that all insoluble elastin content was solubilized, and the assay was performed according to the manufacturer's protocol.

***Biaxial mechanical testing:*** Fresh PA and RV tissues were cut in square $4.5 \times 4.5$ mm$^2$ samples, sprinkled with graphite powder (for displacement tracking), and mounted on a biaxial mechanical tester (BioTester 5000, CellScale, Canada) using a tine sample attachment system (Biorakes with 0.7 mm tine spacing, CellScale, Canada). Samples were submerged in a Dulbecco's phosphate-buffered saline (DPBS) (Gibco, USA, cat# 14190-144) bath at 37 °C and subjected to ten displacement-control preconditioning cycles with displacements of $U_{Max,Long.}$ and $U_{Max,Circ.}$ in the longitudinal and circumferential direction, respectively, generating equal force in two directions (equibiaxial loading condition). Subsequently, the samples underwent nine displacement-control testing protocols with a gradual increase of longitudinal/circumferential load ratios (Figure 1SA and Table 1).

**Table 1.** Prescribed displacement in the longitudinal (Long.) and circumferential (Circ.) direction of pulmonary artery and right ventricle tissue in preconditioning and test cycles.

| Cycles | Displacement in the Long. direction | Displacement in the Circ. direction |
|---|---|---|
| Preconditioning | $U_{Max,Long.}$ | $U_{Max,Circ.}$ |
| Test cycle 1 (P1) | $U_{Max,Long.} + U_{Adj,Long.}$ | ${}^{1}/_{5}\, U_{Max,Circ.}$ |
| Test cycle 2 (P2) | $U_{Max,Long.} + {}^{3}/_{4}\, U_{Adj,Long.}$ | ${}^{2}/_{5}\, U_{Max,Circ.}$ |
| Test cycle 3 (P3) | $U_{Max,Long.} + {}^{1}/_{2}\, U_{Adj,Long.}$ | ${}^{3}/_{5}\, U_{Max,Circ..}$ |
| Test cycle 4 (P4) | $U_{Max,Long.} + {}^{1}/_{4}\, U_{Adj,Long.}$ | ${}^{4}/_{5}\, U_{Max,Circ..}$ |
| Test cycle 5 (P5) | $U_{Max,Long.}$ | $U_{Max,Circ..}$ |
| Test cycle 6 (P6) | ${}^{4}/_{5}\, U_{Max,Long.}$ | $U_{Max,Circ..} + {}^{1}/_{4}\, U_{Adj,Circ..}$ |
| Test cycle 7 (P7) | ${}^{3}/_{5}\, U_{Max,Long.}$ | $U_{Max,Circ..} + {}^{1}/_{2}\, U_{Adj,Circ..}$ |
| Test cycle 8 (P8) | ${}^{2}/_{5}\, U_{Max,Long.}$ | $U_{Max,Circ..} + {}^{3}/_{4}\, U_{Adj,Circ..}$ |
| Test cycle 9 (P9) | ${}^{1}/_{5}\, U_{Max,Long.}$ | $U_{Max,Circ..} + U_{Adj,Circ..}$ |

Force and displacement values in X and Y directions were collected, and the X-Y coordinates of four points on the sample, sprinkled with graphite powder, during the stretch cycles were determined via image tracking.

***Mechanical data processing:*** At neural state, $X_1$ and $X_2$ correspond to the X and Y coordinates in the sample, respectively. During the nine test protocols (P1–P9), the location of the tracked points, determined by image analysis, were mapped to $x_1$ and $x_2$ using $x_1 = \lambda_1 X_1 + F_{12} X_2$ and $x_2 = F_{21} X_1 + \lambda_2 X_2$, where $F_{12}$ and $F_{21}$ are the components of the transformation gradient tensor, and $h/H$ is the sample's deformed/neutral thickness ratio.

Assuming the sample is incompressible and therefore the determinant of the transformation gradient tensor equals 1 (eq. 1), the sample's thickness during deformation was calculated using $h = H/J_{2D}$, where $J_{2D} = \lambda_1 \lambda_2 - F_{12} F_{21}$.

$$\boldsymbol{F} = \partial \boldsymbol{x} / \partial \boldsymbol{X} = \begin{bmatrix} \lambda_1 & F_{12} & 0 \\ F_{21} & \lambda_2 & 0 \\ 0 & 0 & h/H \end{bmatrix} \tag{1}$$

The Green strain tensor was calculated using $\boldsymbol{E} = 1/2\,(\boldsymbol{F}^T \cdot \boldsymbol{F} - \boldsymbol{I})$, and the components of the membrane tension tensor, $\boldsymbol{S}$, were calculated according to:

$$T_{11}^S = \lambda_2 f_1 / (J_{2D} L) \tag{2}$$

$$T_{22}^S = \lambda_1 f_2 / (J_{2D} L) \tag{3}$$

where $f_1$ and $f_2$ are the force readings in the X and Y directions, respectively. $T_{11}^S$and $T_{22}^S$were plotted against $E_{11}$ and $E_{22}$ to represent the experimental membrane tension-Green strain relationships in the longitudinal and circumferential direction of PA and RV tissues. With the assumption of strain energy conservation, the membrane tension and Green strain tensors were related according to:

$$\begin{bmatrix} T_{11}^S & T_{12}^S \\ T_{12}^S & T_{22}^S \end{bmatrix} = \begin{bmatrix} \frac{\partial w}{\partial E_{11}} & \frac{\partial w}{\partial E_{12}} \\ \frac{\partial w}{\partial E_{12}} & \frac{\partial w}{\partial E_{22}} \end{bmatrix} \tag{4}$$

where $w$ is the 2-dimensonal strain energy function. To characterize the mechanical properties of the PA and RV tissues via material constants, a seven-parameter Fung model[1] was used:

$$w = \frac{C_1}{2}(e^Q - 1) \tag{5}$$

where $Q = C_2{E_{11}}^2 + C_3{E_{22}}^2 + 2C_4E_{11}E_{22} + C_5{E_{12}}^2 + 2C_6E_{11}E_{12} + 2C_7E_{22}E_{12}$. Material constants ($C_1$– $C_7$) were determined by minimizing the objective function, $g$, matching the experimental and model-predicted membrane tensions:

$$g = \left\| \left|T^S_{11_{Exp}} - T^S_{11_{Model}}\right| + \left|T^S_{22_{Exp}} - T^S_{22_{Model}}\right| + \left|T^S_{12_{Exp}} - T^S_{12_{Model}}\right| \right\|^2 \tag{6}$$

where

$$T^S_{11_{Exp}} = \lambda_2 f_1/(J_{2D}L) \tag{7}$$

$$T^S_{11_{Model}} = C_1 e^Q (C_2E_{11} + C_4E_{22} + C_6E_{12}) \tag{8}$$

$$T^S_{22_{Exp}} = \lambda_1 f_2/(J_{2D}L) \tag{9}$$

$$T^S_{11_{Model}} = C_1 e^Q (C_3E_{22} + C_4E_{11} + C_7E_{12}) \tag{10}$$

$$T^S_{12_{Exp}} = F_{21} f_1/(J_{2D}L) \tag{11}$$

$$T^S_{12_{Model}} = C_1 e^Q (C_5E_{12} + C_6E_{11} + C_7E_{22}) \tag{12}$$

Among these seven material constants, $C_1$, $C_2$, and $C_3$ were determined to be the most significant and were used to reflect the tissue low-strain stiffness in the longitudinal ($C_1 \cdot C_2$) and circumferential ($C_1 \cdot C_3$) directions (with units of N/m or N/m$^2$ when stiffness is determined based on membrane tension or normalized to the tissue thickness, respectively). $C_2$ and $C_3$ (unitless) governed the slope of the tension-strain curve in the heel and linear (high-strain) regions in the longitudinal and circumferential direction, respectively, and are referred to as high-strain stiffness in the manuscript. The ratio of $C_2/C_3$ was considered as the mechanical anisotropy of tissues. All calculations were performed using a previously developed MATLAB code[2].

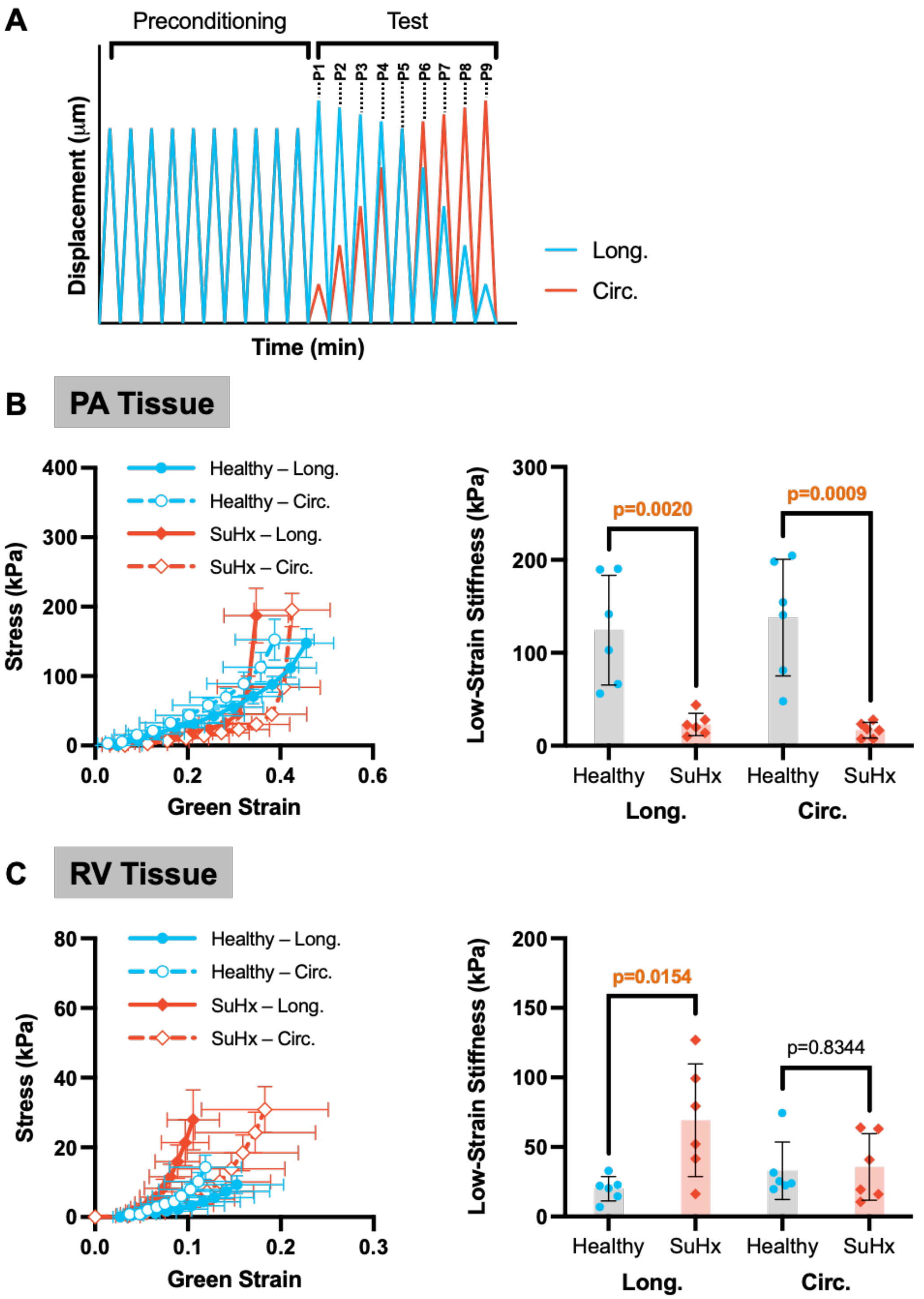


**Figure S1. Representation of pulmonary artery (PA) and right ventricle (RV) tissue mechanics using stress.** **A)** Cyclic loading protocol used to test the mechanical properties of healthy and Sugen hypoxia (SuHx) PA and RV tissues. **B)** Stress-strain behaviour and low-strain stiffness of PA tissue represented in stress in the longitudinal (Long.) and circumferential (Circ.) directions. **C)** Stress-strain behaviour and low-strain stiffness of RV tissue represented in stress. Data are presented as mean ± SD with n=6.

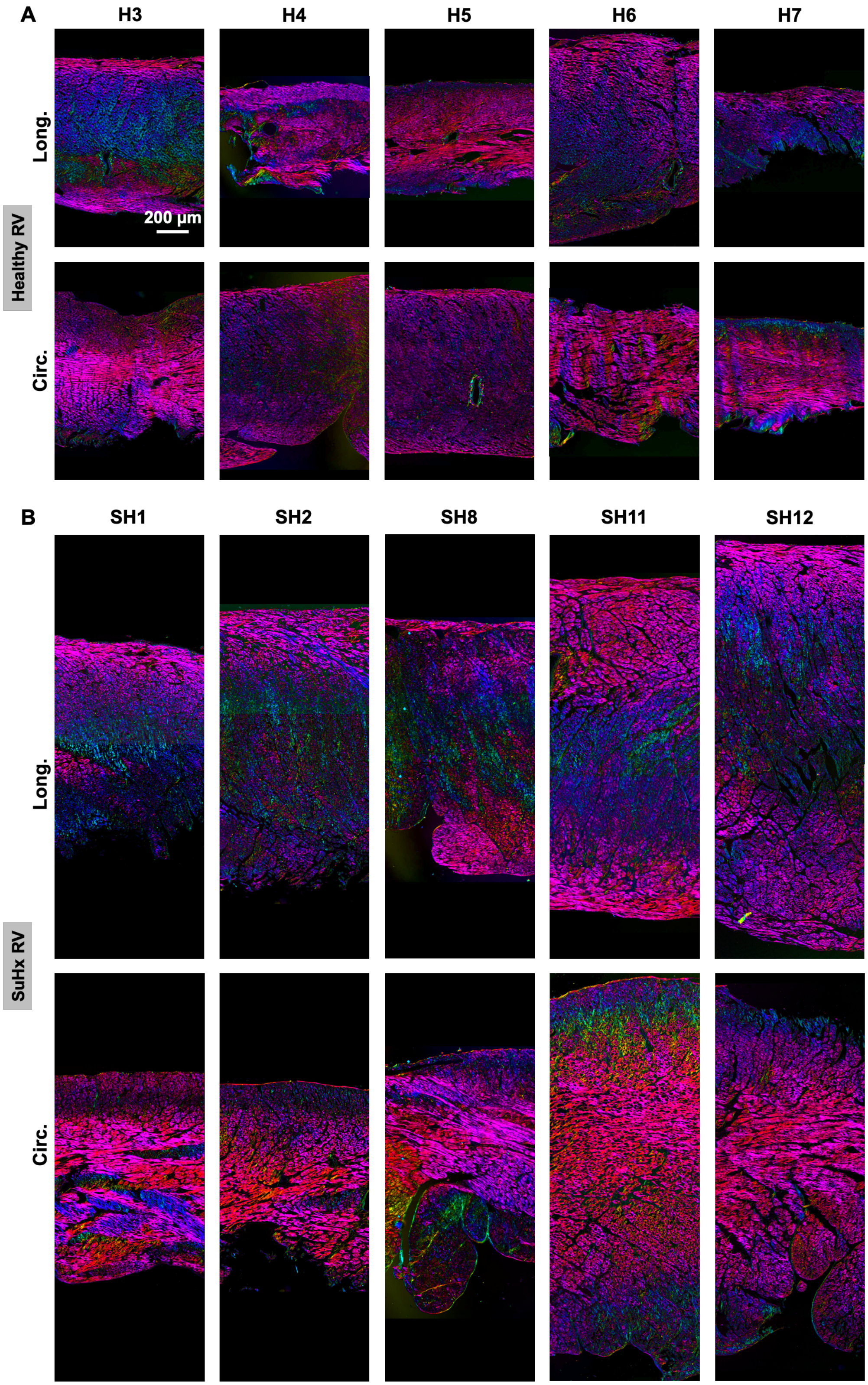


**Figure S2. Polarized light microscopic images of Movat's pentachrome stained sections of healthy and Sugen hypoxia (SuHx) RV tissues in the longitudinal (Long.) and circumferential (Circ.) direction.**

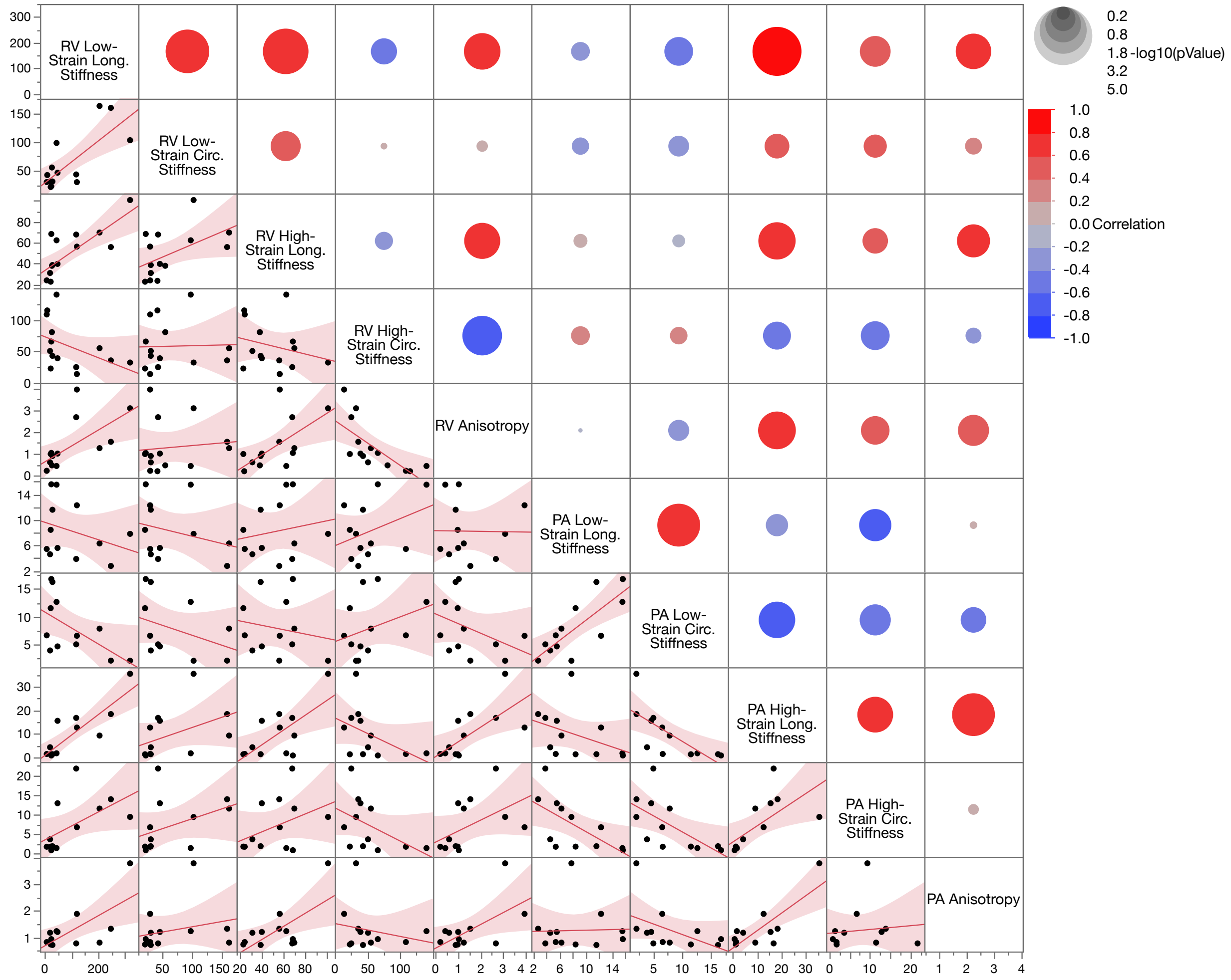


**Figure S3. Correlation between pulmonary artery (PA) and right ventricle (RV) tissue passive mechanics in the longitudinal (Long.) and circumferential (Circ.) directions**. The size and colour of circles show p-value and correlation coefficient, respectively, according to the legends. n=6.

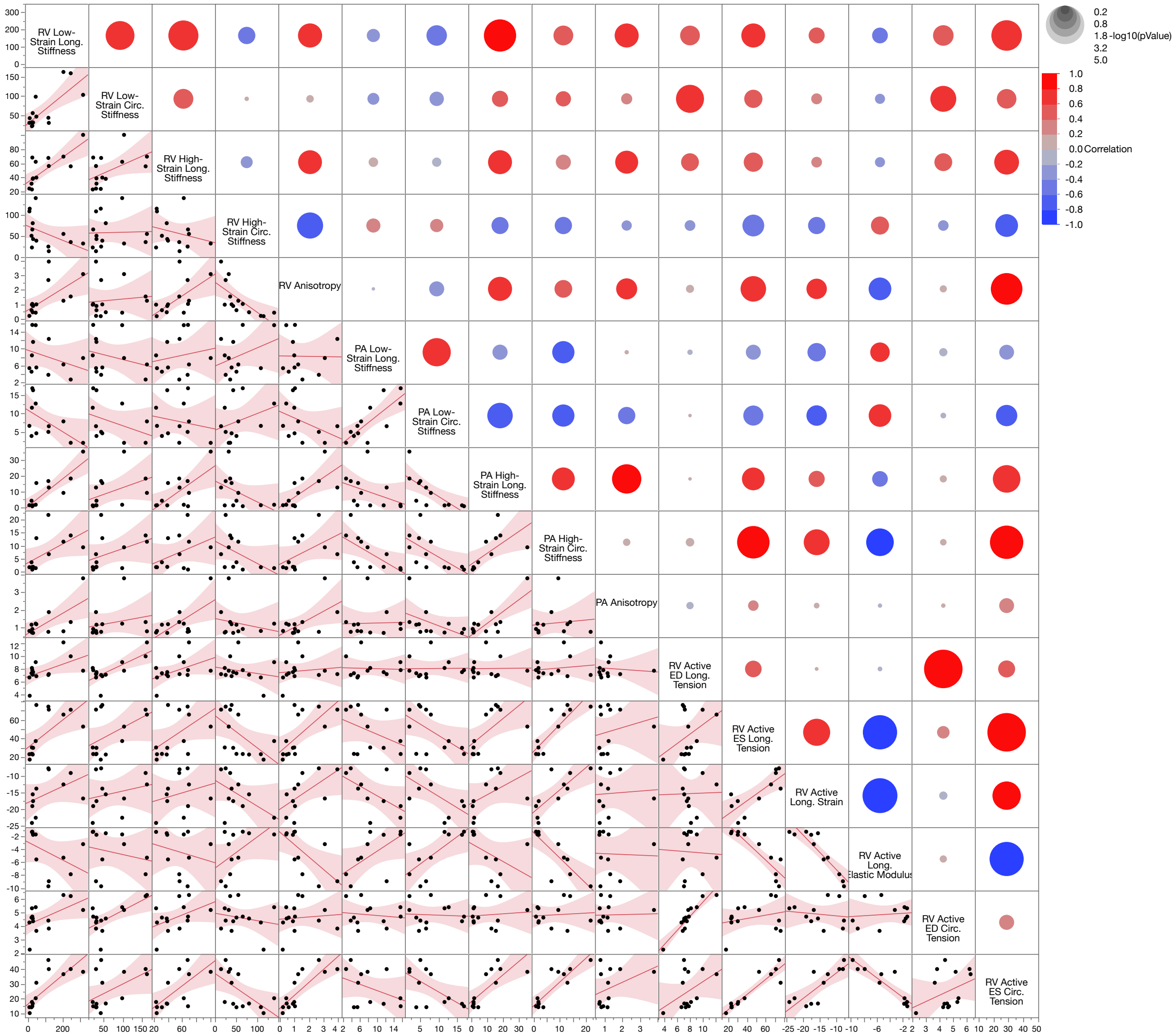


**Figure S4. Correlation between right ventricle (RV) active mechanics and pulmonary artery (PA) and RV passive mechanics in the longitudinal (Long.) and circumferential (Circ.) directions**. The size and colour of circles show p-value and correlation coefficient, respectively, according to the legends. n=6.

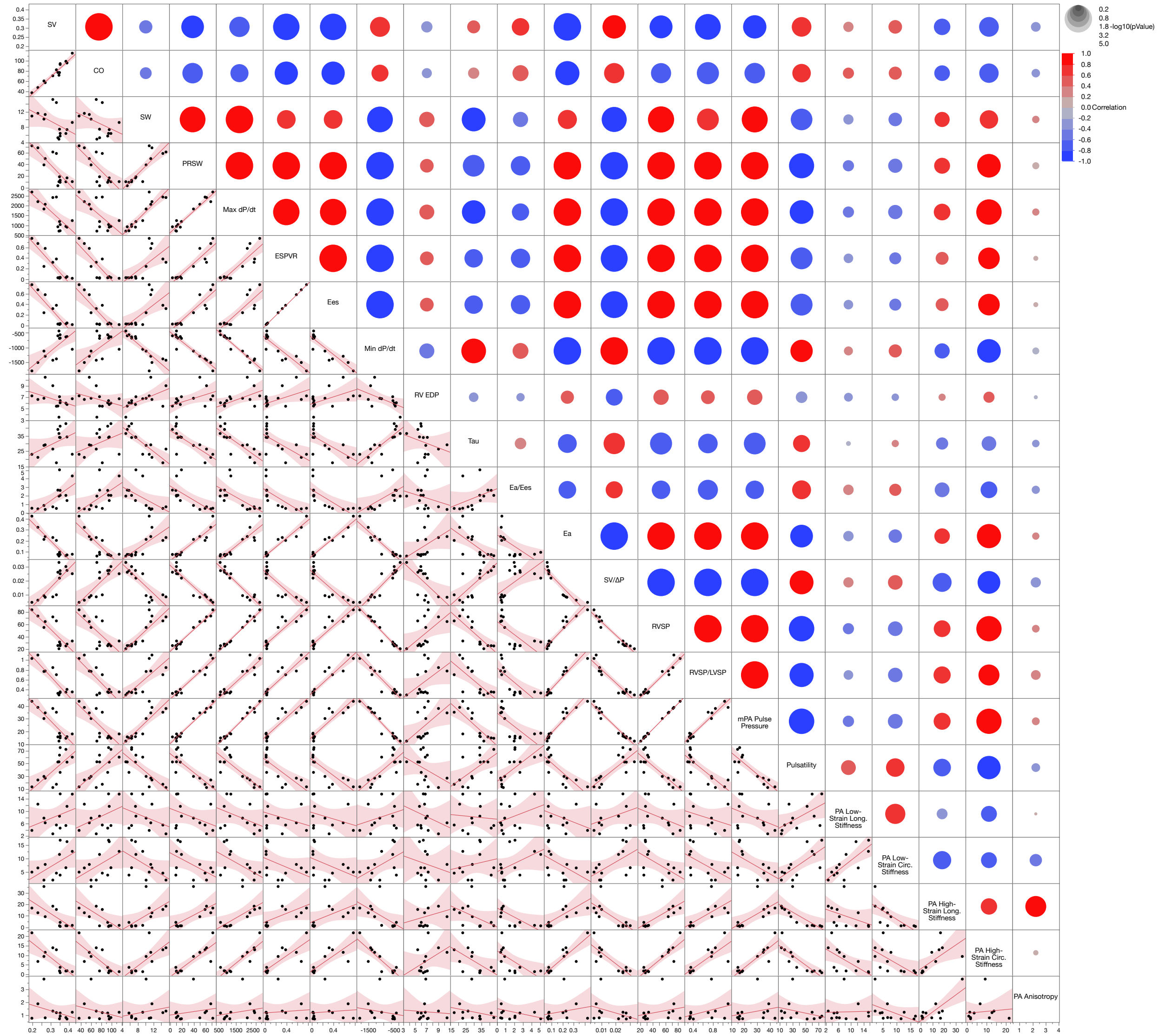


**Figure S5. Correlation between pulmonary artery (PA) tissue mechanics and hemodynamics parameters.** Long.: longitudinal; Circ.: circumferential. The size and colour of circles show p-value and correlation coefficient, respectively, according to the legends. n=6.

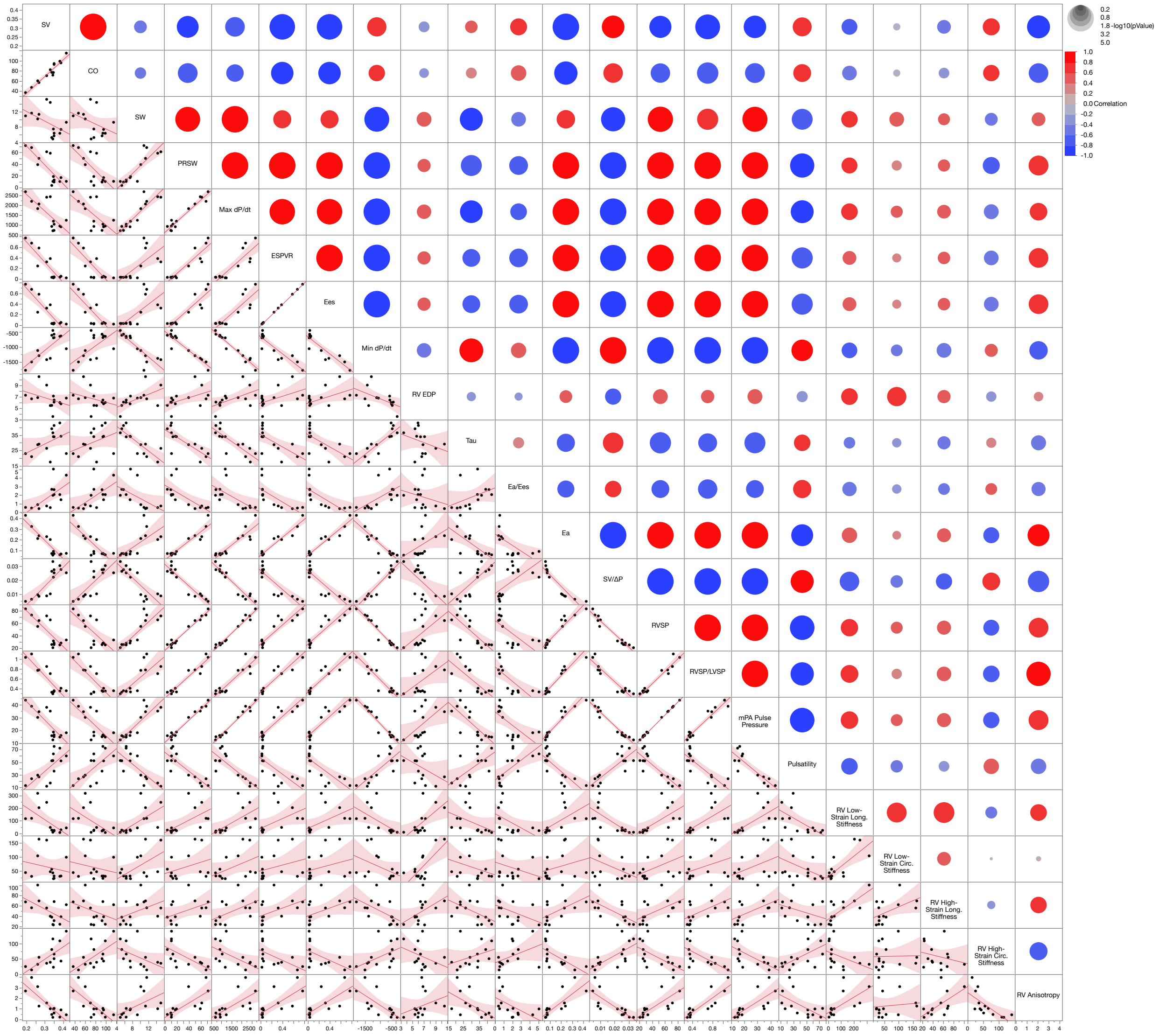


**Figure S 6. Correlation between right ventricle (RV) tissue mechanics and hemodynamics parameters.** The size and colour of circles show p-value and correlation coefficient, respectively, according to the legends. n=6.

### References


1 Sacks, M. A method for planar biaxial mechanical testing that includes in-plane shear. *J Biomech Eng* **121**, 551-555 (1999). https://doi.org/10.1115/1.2835086

2 Labrosse, M., Jafar, R., Ngu, J. & Boodhwani, M. Planar biaxial testing of heart valve cusp replacement biomaterials: Experiments, theory and material constants. *Acta Biomater* **45**, 303-320 (2016). https://doi.org/10.1016/j.actbio.2016.08.036
